# ARGX-119, a therapeutic agonist antibody targeting MuSK

**DOI:** 10.1101/2024.07.18.604166

**Authors:** Roeland Vanhauwaert, Julien Oury, Bernhardt Vankerckhoven, Christophe Steyaert, Stine Marie Jensen, Dana L.E. Vergoossen, Christa Kneip, Leah Santana, Jamie L. Lim, Jaap J. Plomp, Roy Augustinus, Shohei Koide, Christophe Blanchetot, Peter Ulrichts, Maartje G. Huijbers, Karen Silence, Steven J. Burden

## Abstract

ARGX-119 is a novel, humanized, agonist monoclonal SIMPLE Antibody™ specific for muscle-specific kinase (MuSK) that is being developed for treatment of patients with neuromuscular diseases. ARGX-119 is the first monoclonal antibody (mAb) that binds with high affinity to the Frizzled-like domain of human, non-human primate, rat and mouse MuSK, without off-target binding, making it suitable for clinical development. Within the Fc-region, ARGX-119 harbors L234A, L235A mutations to diminish potential immune-activating effector functions. Its mode-of-action is to activate MuSK without interfering with its natural ligand neural Agrin, and cluster acetylcholine receptors (AChRs) in a dose-dependent manner, thereby stabilizing neuromuscular function. In a mouse model for *DOK7* congenital myasthenia (CM), ARGX-119 prevented early postnatal lethality and reversed disease relapse by restoring neuromuscular function and reducing muscle weakness and fatigability in a dose-dependent manner. Pharmacokinetic (PK) studies in non-human primates, rats and mice revealed non-linear PK behavior of ARGX-119, indicative of target-mediated-drug disposition (TMDD) and *in vivo* target engagement. Instability of neuromuscular synapses contributes to symptoms in many neuromuscular diseases for example congenital myasthenia (CM), amyotrophic lateral sclerosis (ALS) and spinal muscular atrophy (SMA). ARGX-119 is a novel, first-in-class MuSK agonist mAb in clinical development. Based on this proof-of-concept study, it has the potential to alleviate neuromuscular diseases hallmarked by impaired neuromuscular synaptic function.

**One sentence summary:** ARGX-119 is a novel first-in-class MuSK agonist monoclonal antibody in clinical development for treatment of neuromuscular diseases.

## Introduction

Diseases of the neuromuscular junction (NMJ) impair neuromuscular transmission, causing debilitating and potentially life-threatening muscle weakness. At the NMJ, motor nerve terminals release the neurotransmitter acetylcholine (ACh), which diffuses across the synaptic cleft and binds to acetylcholine receptors (AChRs) in the muscle membrane. Binding of two ACh molecules opens the ligand-gated ion channel, depolarizing the muscle membrane and initiating an action potential that stimulates muscle contraction [1, 2].

Muscle-specific kinase (MuSK) is a receptor tyrosine kinase essential for the establishment and preservation of the NMJ. MuSK is expressed and required throughout life [1, 3-5], and defects in MuSK signaling are a cause of muscle weakness in congenital myasthenic syndrome (CM) and myasthenia gravis [6]. The transmembrane protein is comprised of an extracellular region with three immunoglobulin-like domains (Ig-like domains 1-3), a Frizzled-like domain (Fz), a transmembrane helix and an intracellular kinase domain [7, 8]. The motor neuron-derived heparan-sulfate proteoglycan neural Agrin induces synaptic differentiation by binding low-density lipoprotein receptor-related protein-4 (LRP4), which promotes association between LRP4 and MuSK and stimulates MuSK tyrosine phosphorylation [9-13]. MuSK phosphorylation leads to recruitment of downstream of kinase 7 (Dok7), essential for anchoring and clustering AChRs on the postsynaptic side of the NMJ, enabling neuromuscular transmission and muscle contraction [7, 14, 15]. MuSK is also required for presynaptic differentiation [3], as MuSK clusters LRP4 which signals back to motor neurons to stimulate differentiation of motor nerve terminals [16].

Agonist antibodies that bind the MuSK Fz-like domain have shown preclinical therapeutic potential in models for Amyotrophic Lateral Sclerosis (ALS) and CM [17-19]. However, these antibodies either recognize proteins in addition to MuSK or fail to recognize human MuSK [17, 20]. Thus, discovery of a MuSK agonist antibody that binds selectively to MuSK in humans and mice would be an important step forward in clinical development of a therapeutic for treating neuromuscular diseases.

The MuSK Fz-like domain is attractive for antibody-targeting, as this domain is dispensable for MuSK function, and agonist antibodies that bind this domain are well tolerated in mice [17-19, 21, 22]. Here, we describe the development of ARGX-119, a monoclonal antibody (mAb) generated by argenx’ proprietary SIMPLE Antibody™ platform, targeting the MuSK Fz-like domain. Binding of ARGX-119 to the MuSK Fz-like domain facilitates dimerization of MuSK, initiating MuSK phosphorylation and inducing postsynaptic differentiation. ARGX-119 is cross-reactive and specific for human, non-human primate (NHP), rat and mouse MuSK. It has a non-linear pharmacokinetic (PK) profile and rescues *Dok7 CM* mice from early postnatal death and disease relapse in adult mice in a dose-dependent manner. These results support the translation of ARGX-119 to be tested in a clinical setting and illustrate the potential of a targeted therapy for treating neuromuscular diseases.

## Results

### Generation of ARGX-119, a MuSK agonist antibody recognizing human, non-human primate, rat and mouse MuSK

ARGX-119 is a humanized antibody derived from llamas that were immunized with the extracellular region of human MuSK (ECD). ARGX-119 bound specifically to the Fz-like domain of human MuSK (Figure 1A). The affinity and cross-species reactivity of ARGX-119 to human, NHP, rat and mouse MuSK was quantified with surface plasmon resonance (SPR) and isothermal titration calorimetry (ITC, Figure 1B,C, Supplemental Figure 2 and Supplemental Table 1), which demonstrated that ARGX-119 binds MuSK across all species tested with similar affinities (KD), and revealed no major pH-dependent dissociation.

**Figure 1:**
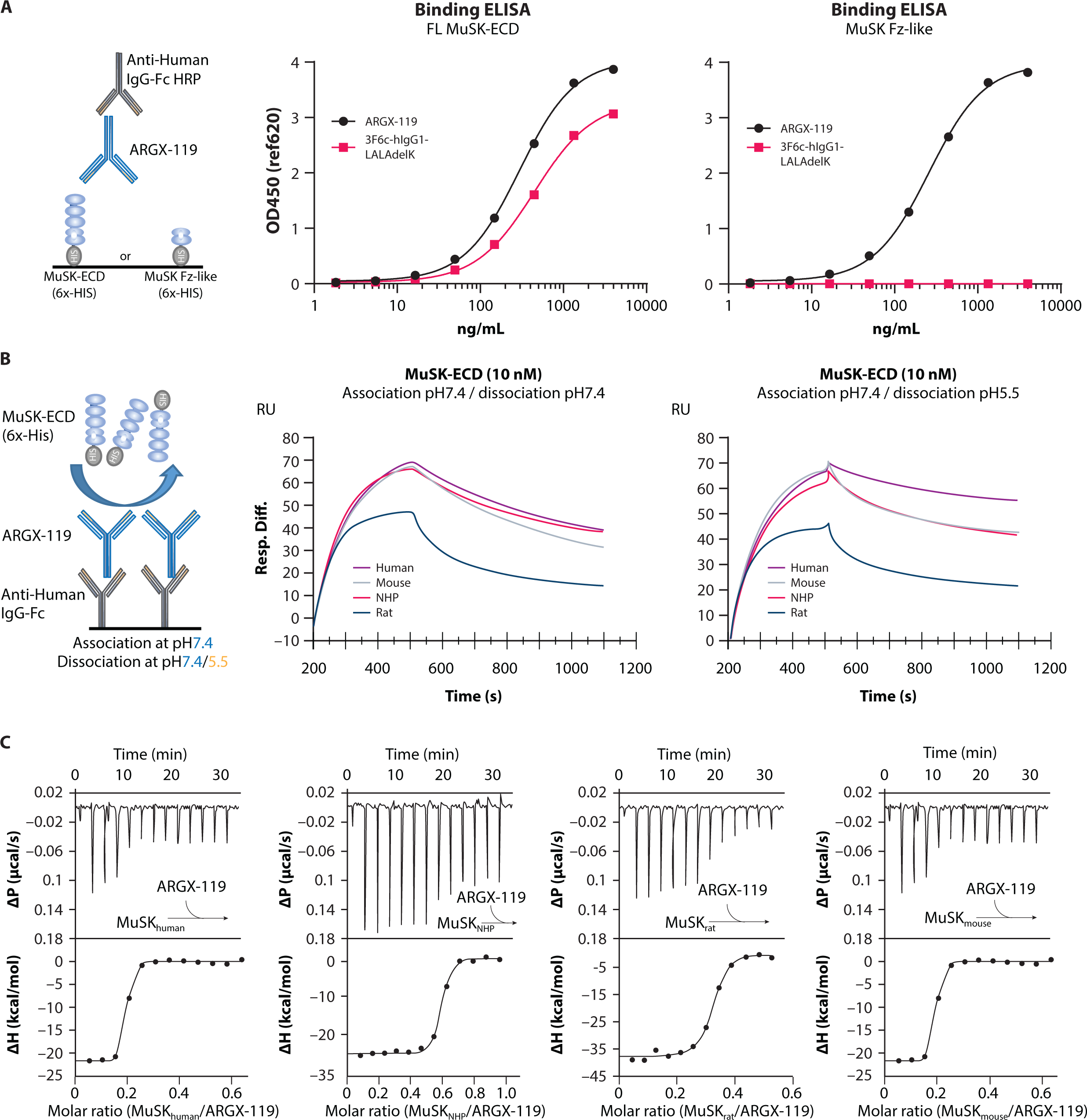
ARGX-119, a MuSK agonist antibody recognizing human, NHP, rat and mouse MuSK. (A) Binding of ARGX-119 or Ig1-like domain binding antibody control (3F6c-hIgG1-LALAdelK) to MuSK-ECD or MuSK Fz-like domain on ELISA. (B) SPR binding of ARGX-119 to MuSK. CM5 chip was coated with anti-human IgG-Fc and ARGX-119 was captured at low densities. Sensorgrams show the binding kinetics of 10 nM human, NHP, mouse and rat MuSK using a dissociation pH of 7.4 or 5.5. Kinetic parameters (C) ITC experiment, investigating the binding thermodynamics of ARGX-119 to human, NHP, mouse and rat MuSK in solution.

ARGX-119 was formatted as a human IgG1 with L234A and L235A (LALA) mutations to reduce binding to Fc gamma receptors (FcγRs) and C1q, diminishing Fc-mediated effector functions such as antibody-dependent cell-mediated cytotoxicity (ADCC) and complement-dependent cytotoxicity (CDC)[23].

Indeed, ARGX-119 had little or no binding to human, NHP, rat and mouse FcγRs. However, we detected a low but measurable affinity to the high affinity IgG-Fc receptor FcγRI (Supplemental Table 2). Similar results were found for binding to C1q (Supplemental Figure 3). These findings indicated that ARGX-119 had the anticipated characteristics; hence, IND (investigational new drug) enabling studies were initiated.

### ARGX-119 stimulates MuSK phosphorylation

We measured the ability of ARGX-119 to stimulate MuSK phosphorylation and AChR clustering in cultured myotubes from different species. ARGX-119 activated the MuSK pathway in a dose-dependent manner across all species tested, including human, NHP, rat and mouse. ARGX-119 was more potent at inducing MuSK phosphorylation on human and NHP myotubes than on rodent myotubes (Figure 2A-F, Supplemental Table 3). Similar results were found for the potency of ARGX-119 to induce AChR clustering (Figure 2G,H and Supplemental Table 3). Analysis of the AChR cluster size distribution revealed that the number and size of AChR clusters were comparable in mouse myotubes treated with Agrin or ARGX-119 (Figure 2I).

**Figure 2:**
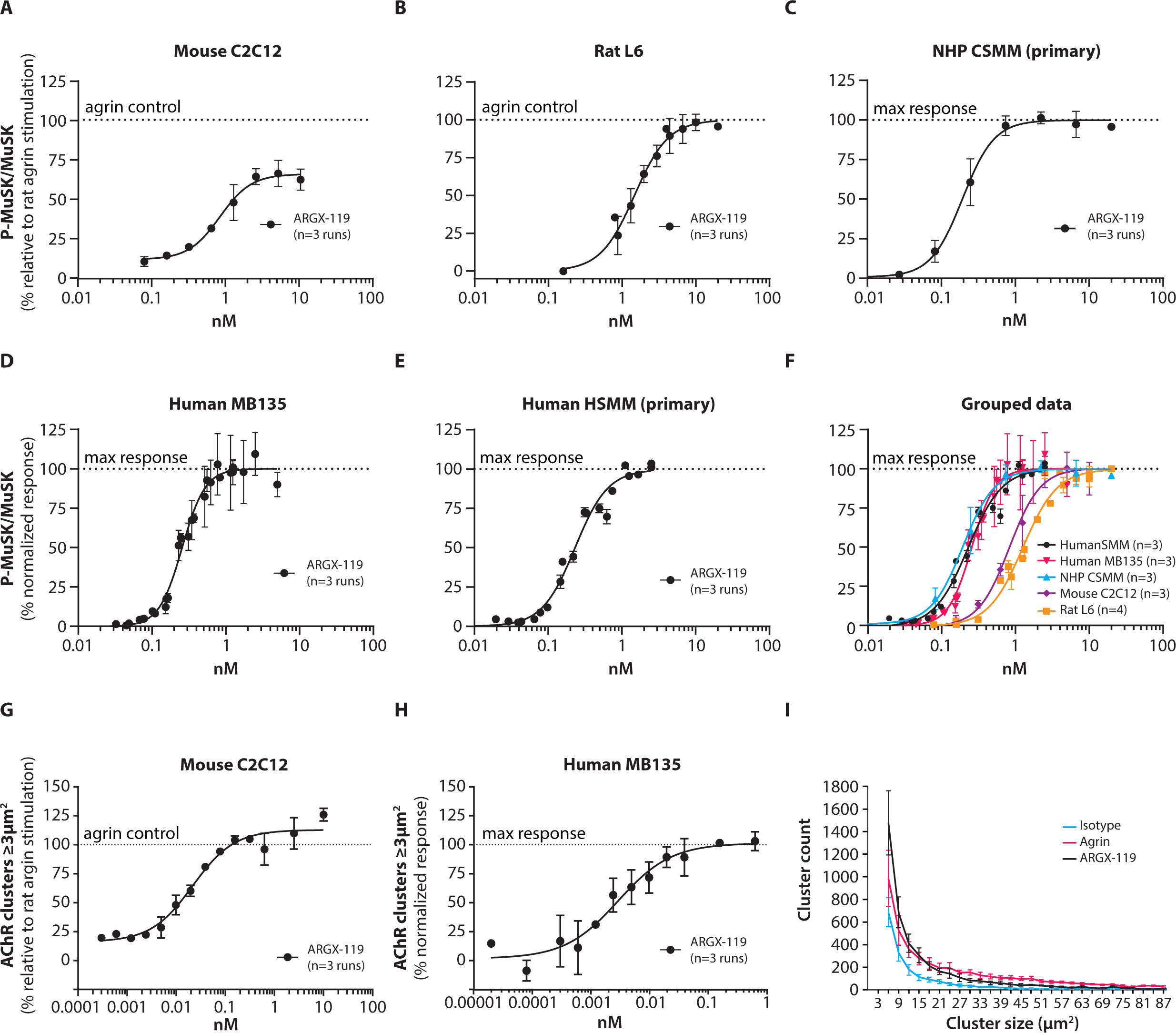
ARGX-119 induces MuSK phosphorylation and AChR clusters *in vitro* across species. MuSK phosphorylation normalized to total MuSK expression in (A) mouse C2C12, (B) rat L6, (C) NHP CSMM primary, (D) human MB135 or (E) human HSMM primary myotubes treated for 30 min (except on mouse C2C12, 10 min) with a dose-range of ARGX-119. Datapoints are the mean ± SD of at least 3 independent duplicates (n=3). (F) The graph shows the grouped data from (A-E). (G) AChR clustering, showing clusters ≥ 3 µm^2^ in mouse C2C12 or (I) human MB135 myotubes, treated for 24 hr with a dose-range of ARGX-119. (I) ARGX-119 and Isotype AChR cluster size distribution normalized to Agrin treatment C2C12 myotubes. AChR clustering data were generated from 3 independent experiments (n=3).

In addition, MuSK phosphorylation in mouse and human myotubes was tested over a wide concentration range of ARGX-119. A bell-shaped dose-response, typical for receptor-dimerizing agonistic Abs [24], was observed over the concentrations tested, which could be rescued by co-incubation of Agrin (Figure 3A-C). Indeed, co-stimulation of ARGX-119 with 0.1 nM Agrin increased MuSK phosphorylation beyond ARGX-119 or 0.1 nM Agrin alone. 0.3 nM ARGX-119 in combination with 0.1 nM Agrin resulted in 100% MuSK phosphorylation, similar to stimulation with 1 nM Agrin (Figure 3D). These findings indicate that ARGX-119 can activate MuSK independently from the natural activator Agrin, leading to increased MuSK phosphorylation.

**Figure 3:**
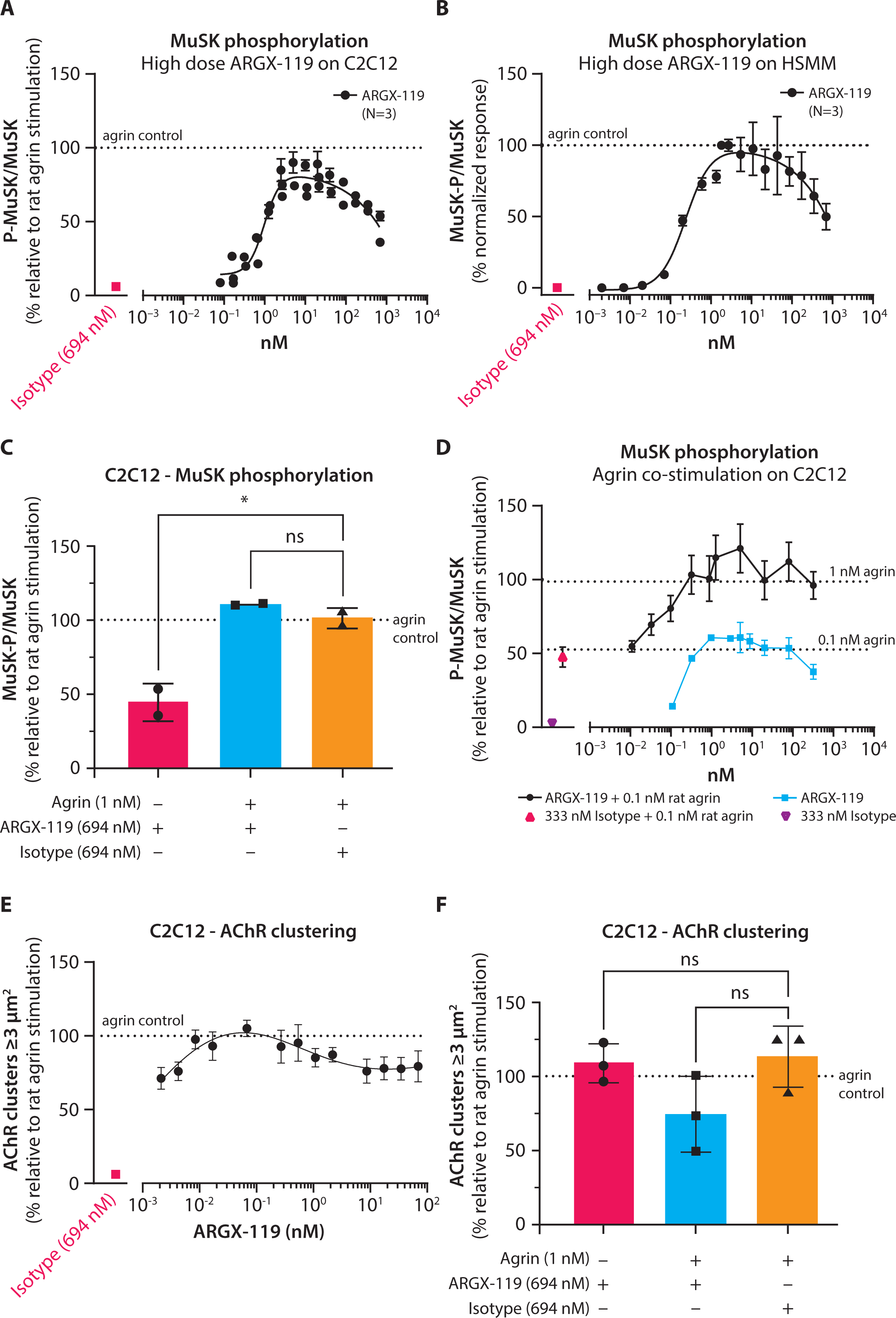
ARGX-119 shows a bell-shaped dose-response and does not interfere with Agrin-induced MuSK phosphorylation and AChR clustering. MuSK phosphorylation normalized to total MuSK expression after a dose-response of ARGX-119 on (A) mouse C2C12 myotubes and (B) human HSMM primary myotubes. Each datapoint represents the mean ± SD of 3 independent experiments (n=3). (C) AChR clustering showing clusters ≥ 3 µm^2^ in mouse C2C12 myotubes treated with ARGX-119. AChR clustering data were generated from 3 independent experiments (n=3). (D) Co-stimulation of ARGX-119 and Agrin on differentiated mouse C2C12 myotubes to induce MuSK phosphorylation. Each datapoint represents the mean ± SD of 2 independent experiments (n=2). The dotted lines represent the 1 nM and 0.1 nM Agrin controls, resulting in 100% and 50% MuSK activation, respectively. (E) MuSK phosphorylation normalized to MuSK expression and Agrin stimulation after treatment with Agrin, ARGX-119, isotype control mAb or in combination on mouse C2C12 myotubes. Each datapoint represents the mean ± SD of 2 independent experiments (n=2). One-way ANOVA with Dunnet’s multiple comparison (ns, not significant; * p<0.05). (F) AChR clustering, showing clusters ≥ 3 µm^2^ in mouse C2C12 myotubes treated with Agrin, ARGX-119, isotype control mAb or in combination. AChR clustering data were generated from 3 independent experiments (n=3). One-way ANOVA with Dunnet’s multiple comparison (ns, not significant).

Next, we investigated whether the reduced MuSK phosphorylation at high concentrations of ARGX-119 (without Agrin) affected AChR clustering. We tested a wide concentration range of ARGX-119 on mouse myotubes and found that ARGX-119 induced AChR clusters (>3µm^2^), at all ARGX-119 concentrations, reaching 80-100% of a saturating dose of neural Agrin (Figure 3E,F, Supplemental Figure 4 shows AChR clusters larger than 15 µm^2^).

These findings suggest that ARGX-119 is a potent activator of MuSK phosphorylation and AChR clustering in human, NHP and rodent myotubes.

### ARGX-119 off-target binding assessment

To assess potential off-target binding of ARGX-119, we used a plasma membrane protein array to measure binding of ARGX-119 to 5474 full-length human plasma membrane proteins and cell surface-tethered human secreted proteins plus 371 human heterodimers [25]. ARGX-119 bound specifically to MuSK and failed to bind other proteins tested. In contrast; a previously described agonist antibody to MuSK, X17, [17] showed off-target binding to two other proteins (EphB1 and EphB2), making it unsuitable for clinical development (Supplemental Table 4).

### ARGX-119 dosing in healthy mice reveals target saturation and target-mediated drug disposition

To determine whether ARGX-119 caused undesired effects in wild-type mice, we repeatedly injected male and female mice intraperitoneally (IP) with ARGX-119 (10 mg/kg at postnatal days (P) 4, P24 and P44) (Figure 4A-G), or twice a week (20 mg/kg) (Supplemental Figure 5), beginning at P4. Chronic dosing of ARGX-119, using either dosing schedule, over two months had no effect on weight gain, motor behavior, the organization of NMJs, and survival when compared to wild-type mice injected with an isotype control Ab.

**Figure 4:**
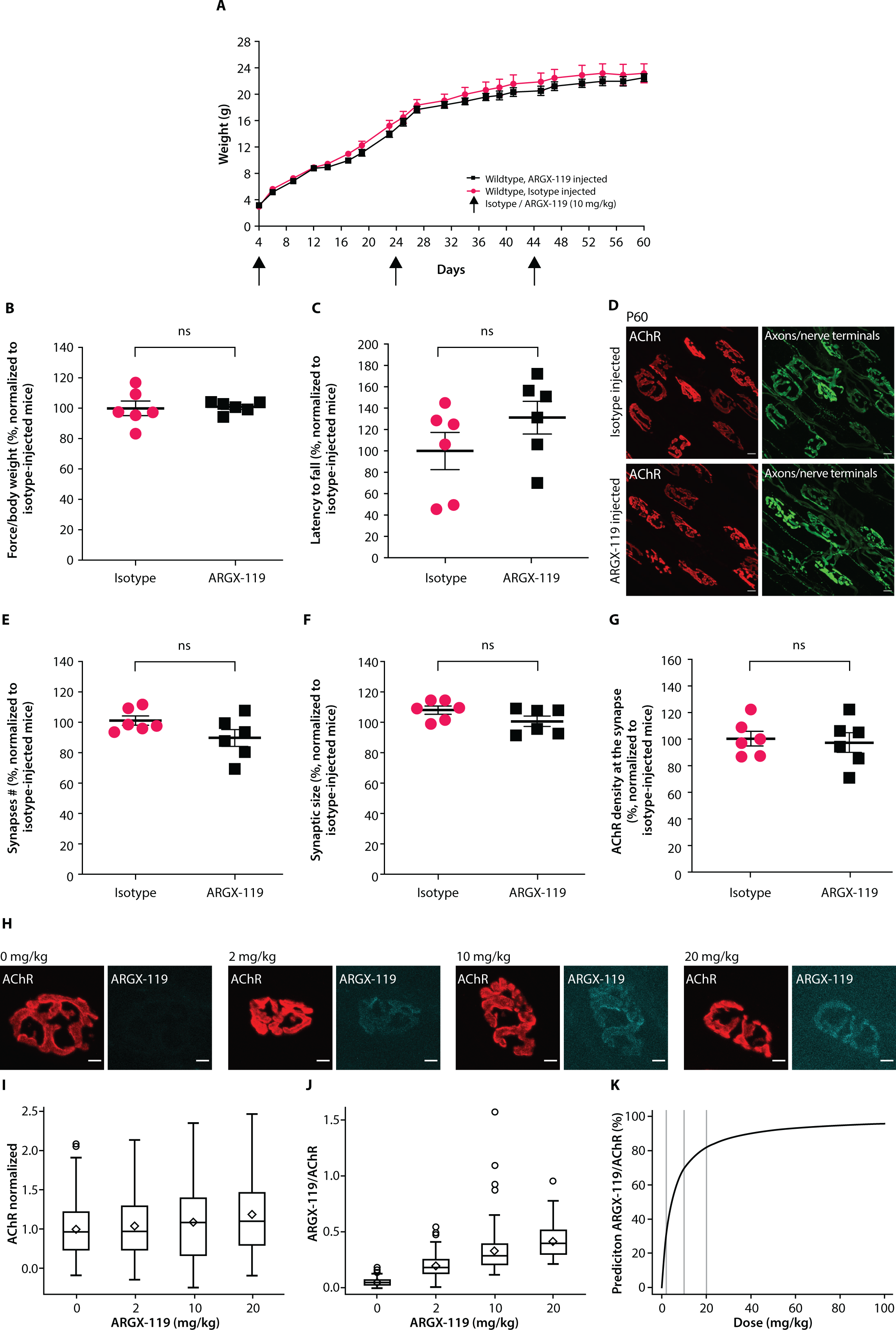
ARGX-119 Is specific for MuSK and binds to MuSK at NMJs in a dose-dependent manner. (A) Wild-type mice injected with ARGX-119 gained weight, like wild-type mice treated with isotype mAb. Plots show averaged data for each group ± SEM. (B-C) Motor performance of wild-type mice injected with ARGX-119, as assessed by grip strength and latency to fall from a rotarod at P60, were similar to isotype-injected wild-type mice. (D) Synapses, from wild-type mice treated with ARGX-119, matured from a simple, plaque-like shape to a complex, pretzel-like shape, characteristic of mature mouse neuromuscular synapses. Scale bar = 10 μm. (E-G) Injection of ARGX-119 in wild-type mice had no effect on synapse number, size or synaptic AChR density. (B-G) Scatter plots show individual data points and means (± SEM) in percentage normalized to isotype-injected mice. Two-sided Student’s *t*-test (ns, not significant). (H) Diaphragm muscles of P30 wild-type mice were stained for AChRs (NMJ) and human IgG (ARGX-119) three days after IP injection of ARGX-119. (I) Box plot representation of normalized AChR intensity per ARGX-119 dosing group. (J) Box plot representation of normalized ARGX-119 / AChR intensity per ARGX-119 dosing group. (K) Graphical representation of the predicted ARGX-119/AChR signal intensities versus ARGX-119 dose (mg/kg).

To gain insight into binding of ARGX-119 to MuSK at the NMJ, healthy wild-type adult mice were injected once IP with 0, 2, 10 or 20 mg/kg ARGX-119. Three days later, we visualized bound ARGX-119 by staining dissected muscle with fluorescently labelled antibodies to human Fc. ARGX-119 bound specifically to NMJs, marked by fluorescently labelled α-bungarotoxin (α-BGT) staining, in the diaphragm muscle in a dose-dependent manner (Figure 4H,J). Statistical modeling based on these datasets estimated that 2.7% of MuSK was bound by ARGX-119 at a dose of 0.125 mg/kg, 53% at 5 mg/kg, 70% at 10 mg/kg and approximately 95% of MuSK at 90 mg/kg ARGX-119 (Figure 4K). AChR expression, as determined by the α-BGT signal intensity, did not significantly change in response to ARGX-119 (Figure 4H-I), indicating that MuSK phosphorylation was maximal at normally innervated NMJs, as described previously [19].

The pharmacokinetics of ARGX-119 were subsequently explored in single-dose studies with either NHP, rats or mice. First, we performed single-dose studies with male and female mice, administered with an IP dose of 0.3, 1.0, 10 or 20 mg/kg. Next, male and female rats were administered once with an intravenous dose (IV, bolus injection) of 0.3, 1.0, 10 or 20 mg/kg. In addition, single-dose studies with male and female NHP, were administered with an IV bolus injection of 0.25, 2.5, 25, or 100 mg/kg. The results of these studies revealed a nonlinear PK behavior in all species, which is indicative of target-mediated-drug disposition (TMDD) (Figure 5A-C). TMDD suggests target engagement and results in more than a dose-proportional increase in overall exposure. The half-life of ARGX-119 in healthy mice at a dose of 10 mg/kg was approximately 2 weeks. No PK differences were observed between male and female animals (Figure 5A-C, Supplemental Table 5).

**Figure 5:**
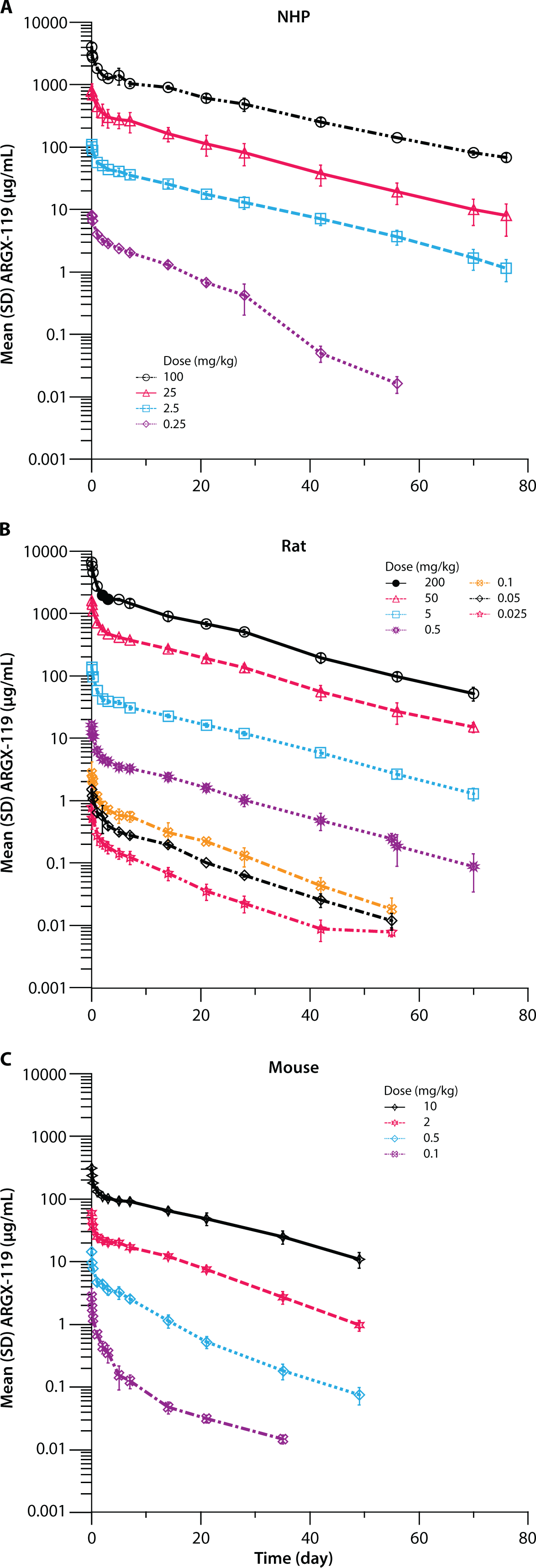
ARGX-119 shows non-linear PK behavior in NHP, rat and mouse. Pharmacokinetic profile of ARGX-119 serum levels over time and doses for (A) NHP, (B) rats and (C) mice.

### ARGX-119 rescues neuromuscular synapse formation, prevents early postnatal lethality in *Dok7* CM mice in a dose-dependent manner

*Dok7* CM mice harbor the most commonly found *DOK7* mutation in patients with *DOK7* congenital myasthenic syndrome (CM), which truncates the protein and leads to impaired MuSK activation [17]. *Dok7* CM mice display multiple deficits consistent with the human disease, including NMJ structural defects, muscle weakness, and fatigability [15, 17]. However, *Dok7* CM mice have a more severe disease presentation than humans with *Dok7* CM and die within 2 weeks after birth.

*Dok7* CM mice were injected with 20 mg/kg ARGX-119 at P4, followed by 10 mg/kg at P18 and P38. All *Dok7* CM mice treated with ARGX-119 (N=10) survived until the end of the study (P60) and had wild-type levels of muscle strength and fatigability at P60, whereas isotype control-treated *Dok7* CM mice (N=11) died on average at P11 (Supplemental Figure 6A-D). The NMJs of the *Dok7* CM mice that were treated with ARGX-119 had the complex pretzel-like shape morphology, which is characteristic of fully mature mouse NMJs, though the number and size of NMJs, as well as the density of synaptic AChRs, were not restored to wild-type levels (Supplemental Figure 6E-H).

To determine the minimal ARGX-119 dose needed to rescue body weight and survival of *Dok7* CM mice, a single dose of ARGX-119 was administered at P4 and body weight and survival were assessed. Eight ARGX-119 doses were tested, ranging from 0.125 mg/kg up to 20 mg/kg (Figure 6A-I). A significant increase in body weight was observed in *Dok7* CM mice that received an ARGX-119 dose of 0.5 mg/kg or higher (Figure 6J). In addition, time to body weight stagnation or loss, and time to disease end point was also significantly lengthened with an ARGX-119 dose of 0.5 mg/kg or higher (Figure 6K, L). These data suggest that a minimal dose of 0.5 mg/kg ARGX-119 at P4 is needed to improve body weight and survival of *Dok7* CM mice.

**Figure 6:**
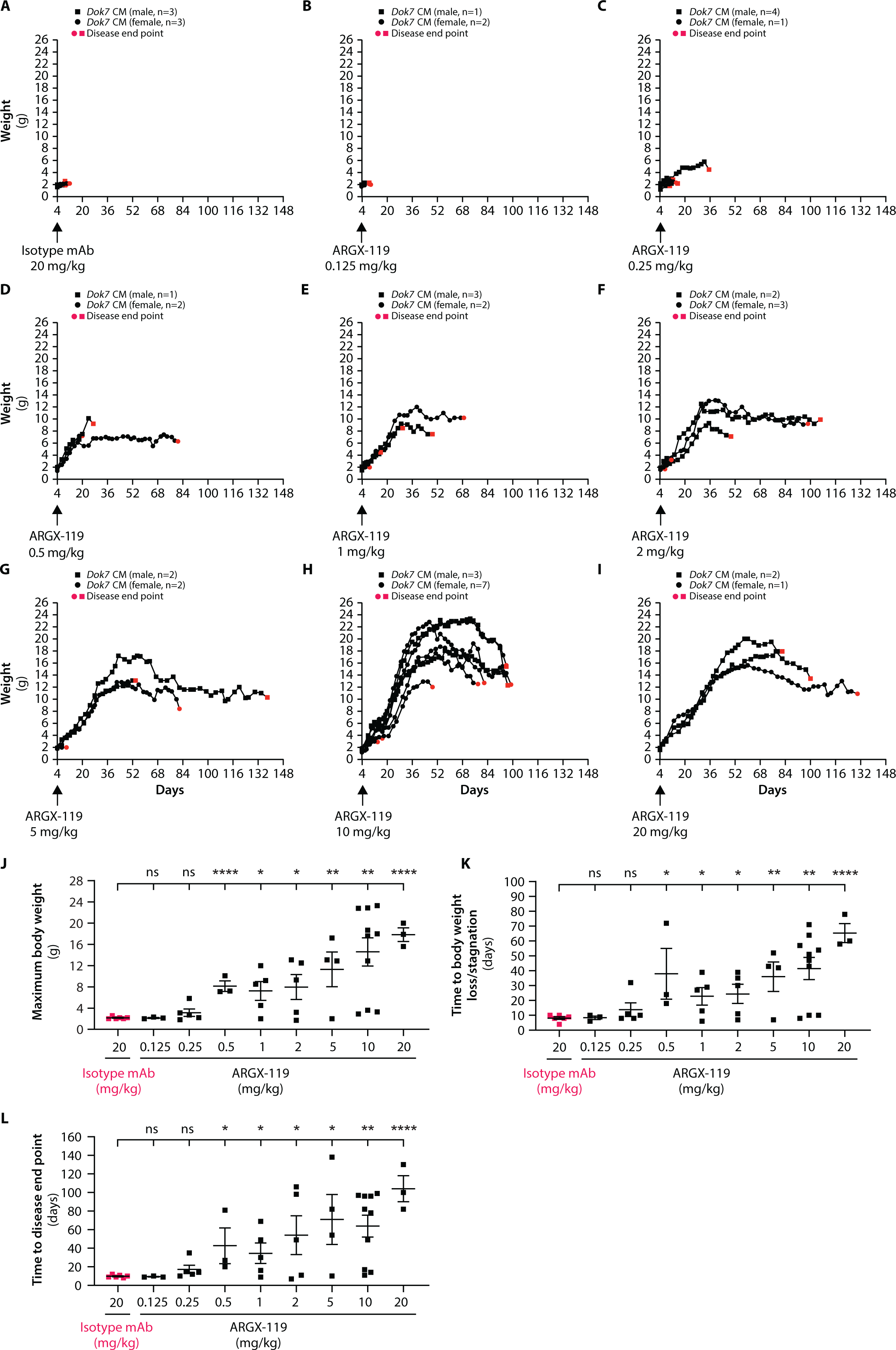
ARGX-119 prevents early postnatal lethality of *Dok7* CM mice in a dose-dependent manner. The body weight and survival of *Dok7* CM mice after a single dose injection at P4 of (A) isotype control at 20 mg/kg or (B-I) ARGX-119 at concentrations ranging from 0.125 mg/kg to 20 mg/kg. All *Dok7* CM mice injected with isotype, or 0.125 mg/kg ARGX-119, and a majority of *Dok7* CM mice injected with 0.25 mg/kg ARGX-119, like untreated mice, did not gain weight and died one to two weeks after birth. Most *Dok7* CM mice injected with ARGX-119 at a dose higher than 0.25 mg/kg gained weight and survived as adults. However, they ultimately began to lose weight and relapsed. Red dots indicate disease end point, when the mice were sacrificed (∼20% weight loss) or died without severe signs of illness. Scatter plots show body weight values for each *Dok7* CM mouse injected with ARGX-119 or isotype. (J) Maximum body weight, (K) time to body weight loss or stagnation and (L) time to death indicate that a single injection of 0.05 mg/kg of ARGX-119 is sufficient to rescue early postnatal death. Scatter plots show individual data points and mean ± SEM values (n≥3 mice for each tested dosing group of isotype or ARGX-119. Two-sided Student’s *t*-test (ns, not significant, p, *<0.05, **<0.005, ****<0.00005).

At ARGX-119 doses of 10 mg/kg and 20 mg/kg, more substantial improvements in body weight and survival times were found. Most mice lived for more than 70 days after a single dose of ARGX-119 (Figure 6H-L). Therefore, a single dose of 10 mg/kg or 20 mg/kg ARGX-119 on P4 was considered fully efficacious for the treatment of *Dok7* CM in mice.

### ARGX-119 reverses disease relapse in adult *Dok7* CM mice in a dose-dependent manner

We regularly monitored the body weight and muscle strength of *Dok7* CM mice injected with ARGX- 119 (10 mg/kg) at P4. At a median of 68 days *Dok7* CM mice began to lose weight and develop signs of muscle weakness. At relapse, the organization of NMJs had deteriorated (Figure 7A-D), and motor performance on a rotarod was diminished (Figure 7E,F). To determine the minimally active dose of ARGX-119 capable of reversing relapse in adult mice, a second dose of ARGX-119 (0.125 - 20 mg/kg) was administered at the time of disease relapse, defined as 5 consecutive days of weight loss. Survival, body weight and muscle function were monitored for 28 days following the second dose (Figure 8A). Relapsed mice treated with ARGX-119 gained weight within days and their motor performance improved for at least four weeks. The minimally active dose range of ARGX-119 was 0.125 to 0.5 mg/kg of ARGX-119. ARGX-119 doses from 2 to 20 mg/kg had a more consistent rescuing effect than lower doses; therefore, ARGX-119 doses of 2 to 20 mg/kg were considered to be fully efficacious in this model (Figure 8B-E and Supplemental Figure 7).

**Figure 7:**
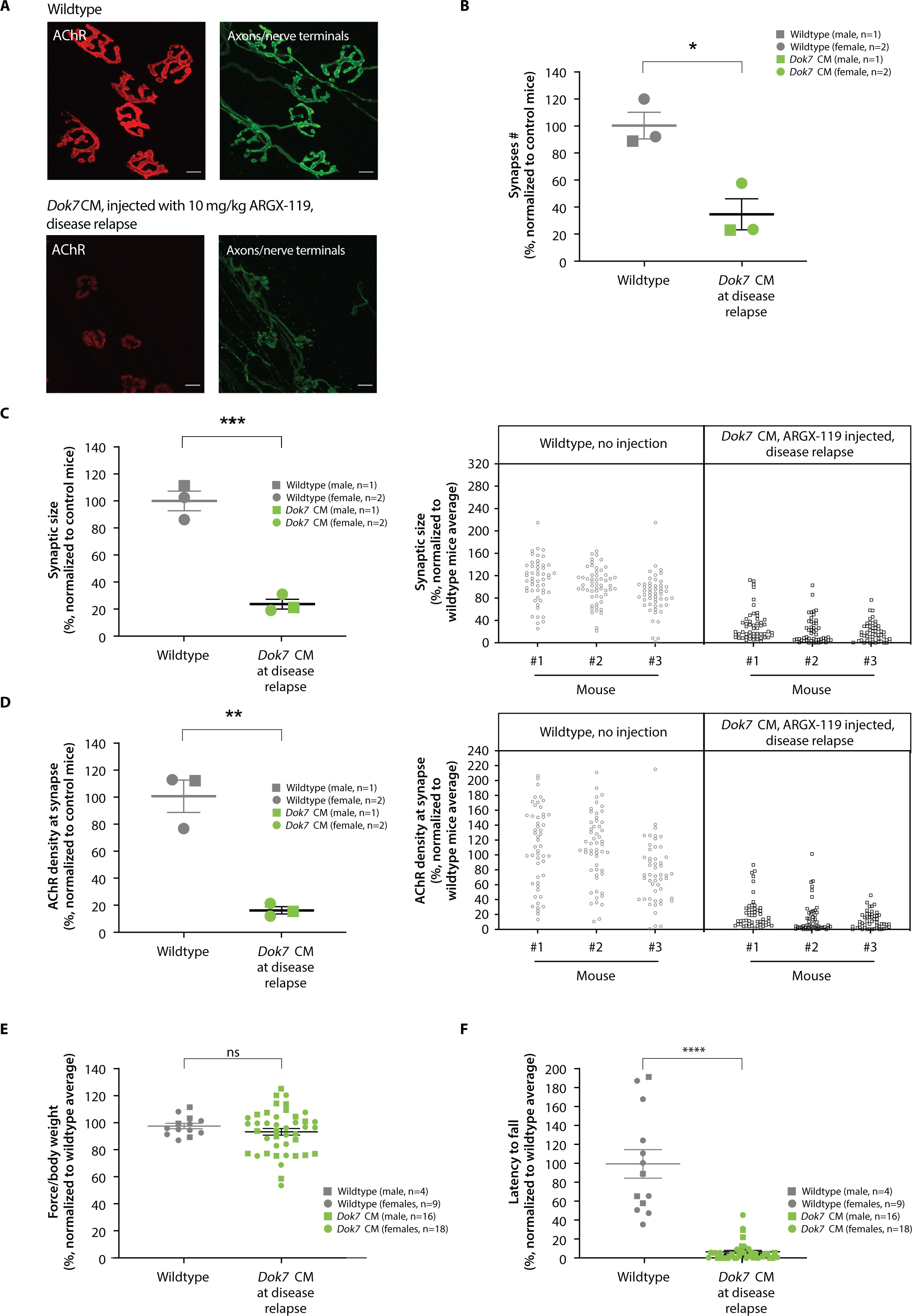
The organization of the NMJ and muscle function deteriorate in *Dok7* CM mice at disease relapse. (A) Diaphragm muscles from wildtype mice (∼P70) and *Dok7* CM mice at disease relapse were stained for AChRs and motor axons/nerve terminals. Scale bar = 10 μm. At disease relapse, synapses number, synaptic size and the density of synaptic AChR were reduced by 65%, 76%, and 84%, respectively, in *Dok7* CM mice compared to wild-type mice. (B, C left panel, D left panel) Scatter plots show the values for each mouse and mean (± SEM) in percentage normalized to wild-type mice. Two-sided Student’s *t*-test (ns, not significant; p, *<0.05, **<0.005, ***<0.0005). (B, C right panel, D right panel). Scatter plots show the percentage values (mean ± SEM) for individual synapses, normalized to wild-type mice. Motor performance of ARGX-119-injected *Dok7* CM mice at disease relapse compared to non-treated wild-type mice at ∼P70. (E-F) Motor performance of wild-type mice (∼P70) and *Dok7* CM mice at disease relapse was assessed by grip strength and latency to fall from a rotating rotarod. *Dok7* CM mice had normal grip strength, but their performance on a rotarod was impaired 10-fold compared to wild-type mice. Scatter plots show the values for each mouse and mean in percentage normalized to wild-type mice average ± SEM. Two-sided Student’s *t*-test (ns, not significant; p, ****<0.00005).

**Figure 8:**
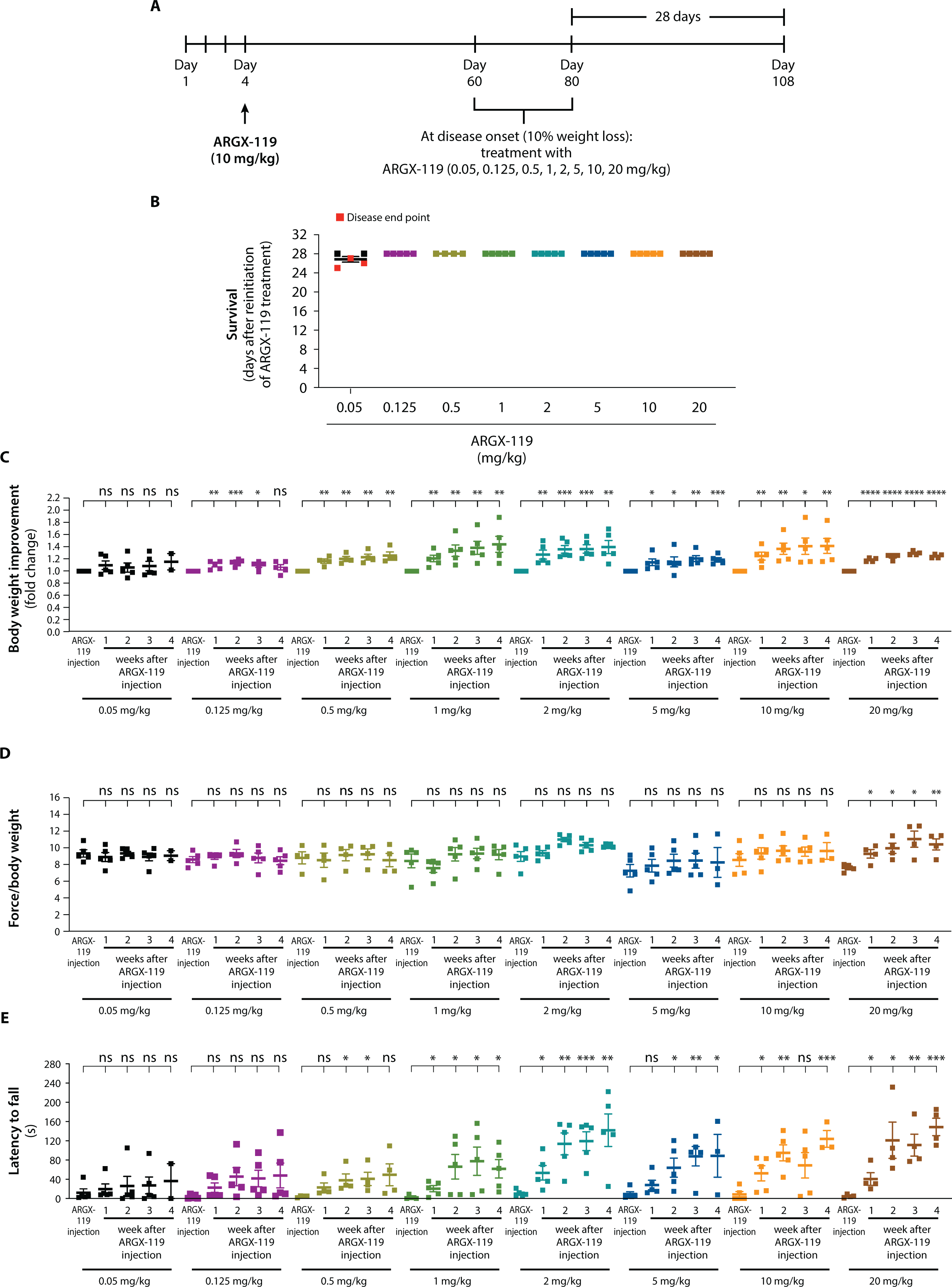
ARGX-119 rescues disease relapse of *Dok7* CM mice in a dose-dependent manner. (A) Strategy to define the minimum dose of ARGX-119 to reverse disease relapse in adult *Dok7* CM mice. (B-C-D-E) When *Dok7* CM mice had lost 10% of their body weight for five consecutive days, they were reinjected with a dose range of ARGX-119 (0.05 to 20 mg/kg) and monitored for 28 days. (B-C) 0.125mg/kg of ARGX-119 was the minimal dose to reverse weight loss and prevent lethality in relapsing adult *Dok7* CM mice. 60% of mice reinjected with 0.05 mg/kg ARGX-119 reached disease end point, whereas 100% of *Dok7* CM mice regained weight by one week and survived to the end of the experiment following injection of ARGX-119 at a dose at 0.125mg/kg or higher. Scatter plots show the values for each *Dok7* CM mouse injected with ARGX-119 and the mean ± SEM. Body weight (fold change) was normalized to body weight on injection day. Two-sided Student’s *t*-test (ns, not significant, p, *<0.05, **<0.005, ***<0.0005, ****<0.00005). (D-E) Reinitiating ARGX-119 treatment at disease relapse improved grip strength and rotarod performances after one week at different doses. (D) 20 mg/kg was the minimal dose to elicit a beneficial effect on grip strength, (E) while only 0.5 mg/kg was needed to ameliorate latency to fall from a rotarod. Scatter plots show the values for each *Dok7* CM mouse injected with ARGX-119 and the mean ± SEM values. Two-sided Student’s *t*-test (ns, not significant, p, *<0.05, **<0.005, ***<0.0005).

Overall, these data suggest that ARGX-119 can rapidly reverse a progressive and lethal phenotype in a therapeutic setting in adult *Dok7* CM mice. Upon signs of body weight loss and muscle weakness, adult *Dok7* CM mice treated with ARGX-119 regained weight within days and their motor performance improved for at least four weeks.

## Discussion

ARGX-119 is a recombinant monoclonal antibody that binds specifically to the Fz-like domain of rodent, NHP (cynomolgus monkey) and human MuSK. The non-clinical data, partially presented here, was supportive for initiation of an ongoing phase 1a clinical trial, investigating the effects of ARGX-119 in healthy volunteers (NCT05670704). The antibody stimulated MuSK kinase activity and AChR clustering in a dose-dependent manner. The agonist antibody was well tolerated in mice, as the organization of neuromuscular synapses, muscle strength, and body weight were unperturbed in mice injected with ARGX-119 and no potential off-target antigens were identified.

ARGX-119 functions as a potent therapeutic in a mouse model of *Dok7* CM, as a single dose of ARGX- 119 prevents early postnatal lethality and rapidly reverses disease relapse in adult *Dok7* CM mice. The prolonged rescue following a single injection of ARGX-119 may be due to the stability of the NMJ once formed, the high safety factor for synaptic transmission at the NMJ, a longer half-life of ARGX-119 in muscle tissue than in blood or a combination of these factors. In any case, the potent and long-lasting benefit from ARGX-119 may allow for chronic dosing of ARGX-119 at monthly or longer intervals.

Together, these findings suggest that ARGX-119 may be a valuable therapeutic for *Dok7* CM and other neuromuscular diseases in humans. In mice, LRP4 expression and MuSK phosphorylation are reduced ∼two-fold during aging, and boosting MuSK phosphorylation by *Dok7* over-expression increases synaptic size and synaptic transmission, leading to greater muscle strength and improved motor performance [26, 27]. Similarly, ARGX-119 has the potential to increase muscle strength and reduce sarcopenia during aging.

Stimulating MuSK, either with agonist antibodies to MuSK or by *Dok7* over-expression has been shown to provide therapeutic benefit in mouse models of amyotrophic lateral sclerosis (ALS), spinal muscular atrophy (SMA), and Emery-Dreifuss muscular dystrophy [18, 19, 22, 28, 29]. Although the precise mechanisms by which boosting MuSK phosphorylation provides benefit in these disease models is not well understood, boosting MuSK phosphorylation stabilizes neuromuscular synapses that would otherwise deteriorate and disassemble in these diseases. Thus, ARGX-119 may provide therapeutic benefit on its own, or in combination with existing treatments, for neuromuscular diseases hallmarked by impaired neuromuscular synapses including ALS and SMA.

## Materials and methods

### Development of ARGX-119

Llamas were immunized with recombinant human MuSK protein. RNA extraction was performed on peripheral blood lymphocytes isolated from the immunized llamas followed by RT-PCR and PCR-cloning strategies to clone the llama Fab sequences in a phagemid as described [30]. Panning phage display selections were performed after one/two or three rounds using either the extracellular domain of human or mouse MuSK or human MuSK lacking the Ig1-like and/or Ig2-like and /or Ig3-like domain (Figure S1). Fz-like domain binding clones were enriched, since this domain is not essential for MuSK function and is conserved across species [21]. Moreover, antibodies against the Fz-like domain cause no obvious harm in mice in contrast to Ig1-like domain binding antibodies [17-19, 31]. Individual clones were screened for binding to human and mouse MuSK by ELISA but revealed poor human/mouse cross-reactive binding. Therefore, light chain shuffling was performed in an attempt to improve species cross-reactivity, and clones binding to both human and mouse MuSK were selected. These Fabs that bound MuSK were produced as human IgG1 antibodies (ImmunoPrecise Antibodies Ltd, Utrecht, the Netherlands), containing the LALA mutations to reduce immune-activating effector functions. The terminal lysine of the heavy chain was removed to decrease heterogeneity. Potency of these antibodies was tested in a MuSK phosphorylation assay on mouse C2C12 myotubes and tested *in vivo* for rescue of early postnatal lethality of *Dok7* CM, mice. Among several agonistic mAbs identified, specific clones were selected for further humanization by germlining of its variable regions using genetic engineering to the framework regions of the closest human germline with the highest identity [30].

### SPR-based kinetic affinity measurements

The affinity of ARGX-119 for human, NHP (cynomolgus monkey), mouse and rat MuSK was measured with a Biacore T200 SPR device (Cytiva). The immobilization buffer was 10 mM sodium acetate, pH 4.5 and the running buffer was HBS-EP+ (10 mmol/L HEPES, 150 mmol/L NaCl, 3 mmol/L EDTA) with 0.05% Tween-20, pH 7.4. Goat anti-human IgG antibodies and the Fcγ fragment specific (109-005-098, Jackson Immunoresearch) were covalently immobilized onto all the flow cells of a Sensor Chip CM5 Series S (Cytiva) at high density using amine coupling chemistry. Next, ARGX-119 was captured and one flow cell treated with running buffer was used as reference. MuSK proteins were then injected over the flow cells using a 2-fold dilution series from 0.31 nM until 20 nM. Dissociation was evaluated using a running buffer at pH 7.4 or pH 5.5. Regeneration after every sample was carried out with acid buffer (10 mM glycine-HCl, pH 2.0). Analysis and calculation of binding data were carried out with Biacore T200 evaluation software version 3.2.

### Binding thermodynamics

The thermodynamic footprint of ARGX-119 binding to the extracellular domain of human, NHP (cynomolgus monkey), mouse and rat MuSK was determined using Isothermal Titration Calorimetry (ITC) on a MicroCal PEAQ-ITC Automated device (Malvern Panalytical). Proteins were thawed once at room temperature (RT) until a small amount of frozen solution remained, before placing the sample on ice to fully thaw. Both MuSK and ARGX-119, were diluted to appropriate working concentrations and dialyzed into 1x PBS pH 7.4, using a Pur-A-Lyzer™ Midi Dialysis Kit (Sigma Aldrich), prior to the start of experiments. Protein concentrations were confirmed using a NanoDrop spectrophotometer. All ITCs were caried out at 37°C using 3 µl injections spaced 150 sec apart, and preceded by a single 0.4 µl injection. Throughout the experiment, a stirring speed of 750 rpm was maintained. For each of the interactions, the affinitie (K_D_), apparent stoichiometries (n), enthalpy change (ΔH), entropy change (ΔS) and the Gibbs free binding energy (ΔG) were determined. Analyses were performed using the MicroCal PEAQ-ITC analysis software (Malvern Panalytical).

Data were analyzed using NITPIC version 1.3.092,93 using the default parameters. Calculated heats and error estimates of all injections were transferred to Sedphat version 1.294. Interactions were modeled using the “A + B + B < - > {AB} + B < - > ABB with 2 symmetric sites, macroscop K” model with the following global parameters: incfA = incfB = 0, not refined; Log(Ka1) = 6, refined; dHAB = −10, refined; Log10(Ka2/Ka1) macroscopic = −0.6, not refined; dH(AB)B-dHAB=0, not refined. Under “experiment parameters”, it was allowed to fit a baseline and estimate a local correction factor for the cell concentration. Buffer, pH and temperature were completed and, given our setup, we selected “titrate A into B”. After a global fit, the estimated thermodynamic parameters of the ABB reaction were calculated from those of the AB estimates. GUSSI version 1.4.2 was used to generate a figure containing the thermogram and isotherms95. Representative ITC thermograms were selected and overlaid with the calculated thermodynamic parameters of the interactions.

### MuSK-binding ELISA

Recombinant human MuSK ECD or the MuSK Fz-like domain was coated overnight at 4°C on a MaxiSorp plate (Thermo Scientific). The next day the assay plate was blocked with 1% casein blocking buffer (Bio-Rad) for 1 hr at RT. ARGX-119 and MuSK Ab 3F6c-hIgG1-LALAdelK (a MuSK MG patient derived MuSK monoclonal targeting the Ig-like 1 domain of MuSK [32]) test dilutions were prepared fresh by diluting the stock solution in assay buffer (0.1% Casein in 1xPBS). These antibody dilutions were added in duplicate to the assay plate and incubated for 2 hr at RT. ARGX-119 was detected using a HRP-conjugated goat F(ab’)2 anti-human IgG-Fc polyclonal antibody (Abcam) for 1h at RT followed- by addition of the TMB substrate. Color development reaction was terminated by adding sulfuric acid, and OD 450nm was measured on an Infinite M Nano plate reader (Tecan). Results were processed using GraphPad Prism 9.0 software.

### MuSK phosphorylation assay

*MuSK stimulation.* MuSK phosphorylation was measured in myotubes. Myotubes were treated with ARGX-119 and Agrin with the indicated concentration for 30 minutes (min), unless stated otherwise. Recombinant neural rat Agrin (R&D Systems) was included as a positive control using a saturating concentration of 1 nM. The isotype control (Mota-hIgG1-LALAdelK), was used at the same concentration as the highest applied concentration of ARGX-119. During the ‘high dose’ experiments, ARGX-119 was serially diluted starting from as high as 694 nM. For the co-stimulation experiments of ARGX-119 with rat neural Agrin, a non-saturating concentration of 0.1 nM Agrin, in combination with serial dilutions of ARGX-119 starting from 333 nM, was used. Cells were detached and lysed using the lysis buffer from a human phosphotyrosine MuSK ELISA kit (Item J, RayBiotech) supplemented with 2 mM activated sodium orthovanadate (Na3VO4) and protease and phosphatase inhibitor cocktails (Roche). Lysates could be stored at -80°C, but MuSK extraction was routinely immunoprecipitated immediately.

*Extraction and detection of P-MuSK/MuSK on MSD.* Immunoprecipitation of MuSK was initiated by adding biotinylated MuSK-Ig1 Ab 3F6c-hIgG1-LALAdelK to each of the lysates followed by an overnight incubation at 4°C. Bound antigen-antibody complexes were precipitated using streptavidin-coated magnetic beads (Promega) for at least 1 hr at 4°C, and the beads were washed extensively with lysis buffer. MuSK proteins were eluted from the streptavidin-coated magnetic beads using 320 mM acetic acid (pH 2.6-2.7) followed by a neutralization step using 1 M Tris (pH 9.5).

Simultaneously, a streptavidin-coated MSD plate (Meso Scale Discovery) was blocked, and biotinylated MuSK-Ig1 Ab 3B5-hIgG4 was captured on the plate. MuSK, eluted from the streptavidin- coated magnetic beads, was loaded in quadruplicate on the plate, which allowed for detection of both total MuSK (via incubation with a mix of polyclonal MuSK Abs (PA1-1741) (Thermo Scientific and MBS9205728, MyBioSource) and tyrosine phosphorylated MuSK (via incubation with a mix of mouse anti-phosphotyrosine clone 4G10 (05-321 Millipore Corp) and clone PY20 (ab10321, Abcam) in the same sample in duplicate. Final detection occurred by applying SULFO-TAG conjugated antibodies, respectively anti-rabbit IgG for total MuSK detection (32AB-1, Meso Scale Discovery) and anti-mouse IgG for phosphorylated MuSK detection (R32AC-1, Meso Scale Discovery). Bound antibodies were detected via electrochemiluminescence (ECL) using the Quickplex MSD (SQ 120 Meso Scale Discovery).

*Analysis of P-MuSK/MuSK data.* In order to compare datasets from different experiments performed on different days, a normalization step was performed. The levels of ARGX-119-induced MuSK phosphorylation were normalized to the Agrin-treated conditions for mouse C2C12 and rat L6 myotubes. For reasons not fully understood, perhaps due to differences in LRP4 expression, neural rat Agrin and neural human Agrin (in-house produced, data not shown) did not potently induce MuSK phosphorylation on human and NHP (cynomolgus monkey) myotubes. These datasets were further normalized by the following formula: normalized response (%) = 100*((response – bottom)/(top – bottom)), where the bottom represents the isotype-treated signal and the top represents the maximal signal. For the titration experiments, the normalized data were fit to a four-parameter non-linear regression model using GraphPad Prism 9.0 from which the EC5, EC10, EC25, EC50, EC75 and EC95 concentrations were calculated. For the ‘high dose’ experiments, data were fit to a bell-shaped non-linear regression model and a one-way ANOVA with Dunnett’s multiple comparisons test was conducted to determine statistical significance using GraphPad Prism 9.0 software. Finally, for the Agrin co-stimulation experiments, data was normalized to the data for 1 nM Agrin and calculated using Graphpad Prism 9.0.

### Stimulation and detection of AChR clusters

*MuSK stimulation.* Mature muscle myotubes had to be formed before AChR clustering experiments were initiated. All stimulations occurred for 24 hr unless stated otherwise. The stimulation times and test concentration range of ARGX-119 was optimized per cell line. Recombinant neural rat Agrin (R&D Systems) was always included as a positive control using a saturating concentration of 0.11 nM. The isotype control, Mota-hIgG1-LALAdelK, was always used at highest applied concentration of ARGX-119. During the ‘high dose’ experiments, ARGX-119 was serially diluted starting from as high as 694 nM with and without co-stimulation of 1 nM Agrin.

*Detection of AChR clusters.* After 24 hr, mouse C2C12 myotubes were stained with Alexa Fluor^TM^ 488- conjugated alpha-Bungarotoxin (α−BGT) (B13422, Thermo Fisher Scientific) and Hoechst (62249, Thermo Fisher Scientific) for 30 min and subsequently fixed with 4% paraformaldehyde (PFA, JT Baker), washed and stored in PBS at 4°C. Human MB135 cells were fixed with 4% PFA for 24 hr and sequentially stained with mAb35-mIgG2c, produced and purified at Evitria SA, which binds AChRs [33], followed by a goat anti-mouse IgG secondary antibody conjugated to Alexa Fluor^TM^ 488 (A32723, Invitrogen). Simultaneously with AChR staining, nuclei were stained with Hoechst, and the cells were washed and stored in PBS at 4°C. The plates containing the cells could be stored at 4°C for up to a week, but the plates were routinely analyzed within 24 hr. Imaging of AChR clusters was performed with a AF6000 fluorescence microscope (Leica AF6000) using a 20x objective. Five visual fields per well and four wells per condition were selected, based on evenly spread and clearly visible mature myotubes in the brightfield channel.

*Analysis of AChR clusters.* Analysis of AChR clustering was performed using ImageJ (1.52n). We selected for AChR clusters in mouse C2C12 and human MB135 myotubes that were at least 3 μm^2^ in size. ARGX-119-induced AChR clustering on mouse C2C12 myotubes were normalized to the levels induced by rat neural Agrin. Since recombinant rat neural Agrin was not potent on MB135 human myotubes, we normalized the data as a expressed response (%) = 100*((response – bottom)/(top – bottom)), where the bottom represents the isotype treated signal and the top represents the maximal signal. A one-way ANOVA with Dunnett’s multiple comparisons test was used to determine statistical significance and was conducted using GraphPad Prism 9.0 software.

### Off-target binding

Off-target binding interactions of ARGX-119 and X17 [17] were assessed using the human plasma membrane protein cell array platform (Retrogenix Ltd, Chinley, United Kingdom). The antibodies were screened for binding against fixed human HEK293 cells, individually expressing 5475 full-length human plasma membrane proteins and cell surface-tethered human secreted proteins plus 371 human heterodimers. Each library hit was re-expressed, along with 2 control receptors, and re-tested with 1 µg/mL of test antibody or control treatments. This was performed both on fixed cells and on live cells in the absence of fixation. First, Pre-screens were undertaken to determine the level of background binding of the test antibody to untransfected HEK293 and cells over-expressing MuSK. These data were used to assess the suitability and optimal concentration for onward screening. Second, in the library screen, the test antibody was screened for binding against fixed HEK293 cells over-expressing 5475 individual full-length human plasma membrane proteins and cell surface-tethered human secreted proteins, as well as a further 371 human heterodimers. Finally, in the Confirmation/Specificity screens, all library hits were re-expressed, and probed with the test antibody or control treatments, to determine which hit(s), if any, were repeatable and specific to the test antibody. This was performed both on fixed and live cells.

### Evaluation of wild-type C57BL/6-CBA mice

*Injection of antibodies in C57BL/6-CBA mice.* ARGX-119 was evaluated in healthy, C57BL/6-CBA mice. Mice received either 3 IP doses of 10 mg/kg at P4, P24 and P44 of ARGX-119 (n=3 males, n=3 females) or an isotype control mAb (n=3 males, n=3 females). Alternatively, mice were IP dosed twice per week, beginning at P5, with 20 mg/kg ARGX-119 or an isotype control mAb IP, until 60 days of age.

*Body weight, muscle strength and motor coordination measurements.* Body weight was determined 2x per week using a Ohaus PX2202/E Pioneer Precision Balance. Grip strength of male and female mice was measured using a grip strength apparatus (Bioseb, BIO-GS3) at P60 as described [17]. To measure all-limb grip strength, mice were positioned in the center of a metal grid and held gently at the base of their tail. Mice were allowed to grip the grid with both forelimbs and hindlimbs, and while mice were pulled back steadily, until mice lost grip with the grid. The grip strength meter digitally displayed the maximum force applied (in grams) as the grasp was released. The means of six consecutive trials of all-limb measurements for each mouse were taken as an index of all-limb grip strength. Mice were given an interval of 10-15 sec between trials. To enhance the robustness and reliability of the grip strength assessment, all measurements were taken by the same experimenter. Motor coordination of male and female mice at P60 was assessed on a rotarod (AccuRotor four-channel, Omnitech Electronics, Inc.). Mice were placed on the rotarod (3.0-cm rotating cylinder) rotating at 2.5 rpm, and either the speed of rotation was increased linearly to 40 rpm over the course of 5 min or speed of rotation was constant at 300 rpm. The time to fall from the rod was measured. Mice were subjected to three trials at 5 min intervals and the longest latency to fall from the three trials was recorded. The two rotarod paradigms were done on the same days with 1 hr time interval between paradigms. A student t-test was used to determine statistical significance and was analyzed using GraphPad Prism 9 software.

*Immunohistological analysis of diaphragm NMJs.* Diaphragm muscles were dissected from P60 mice in oxygenated L-15 medium. The muscles were pinned onto Sylgard-coated dissection dishes, fixed for 1.5 h in 1% PFA and blocked for 1 h in PBS with 3% BSA (Sigma IgG free) and 0.5% Triton X-100 (PBT). Diaphragm muscles were stained with Alexa 488-conjugated α-BGT (Invitrogen) to label AChRs and with antibodies against β-TUBIII (Synaptic Systems 302302) or Synapsin 1/2 (Synaptic Systems, 106002) to label motor axons and nerve terminals, respectively. The antibodies were force-pipetted into the muscle, and the muscles were incubated overnight at 4 °C on an orbital shaker in a humidified chamber. Diaphragm muscles were washed 10 times over the course of 5 h with PT (PBS with Triton X-100) at RT and rinsed in PBS before the muscle was whole-mounted in 50% glycerol. Images were acquired with a Zeiss LSM 800 confocal microscope using ZEN software. Adjustments to detector gain and laser intensity were made to avoid saturation. The number and size of synapses, as well as the density of synaptic AChRs were quantified using FIJI/ImageJ software, as described [34]. More than 50 synapses per diaphragm muscle from each mouse were quantified. A two-sided Student’s t-test was used to determine statistical significance and was conducted using GraphPad Prism 9.0

### Target engagement of ARGX-119 in C57BL/6-CBA mice

*Injection of ARGX-119 in C57BL/6-CBA mice.* Healthy C57BL/6-CBA mice at postnatal day 30 were injected IP with different doses (0, 2, 10 or 20 mg/kg) of ARGX-119. ARGX-119 was diluted in PBS prior to injection.

*Immunohistological analysis of diaphragm NMJs.* Three days after injection of ARGX-119, mice were sacrificed, and diaphragm muscles were dissected in oxygenated L-15 medium. The muscles were pinned onto Sylgard-coated dissection dishes, fixed for 1.5 h in 1% PFA and blocked for 1 h in PBS with 3% BSA (Sigma IgG free) and 0.5% Triton X-100 (PBT). Diaphragm muscles were stained with Alexa 488-α-BGT to label AChRs (red) and Alexa 647 goat anti-human IgG, F(ab’)₂ fragment specific to label ARGX-119 (cyan). ARGX-119 has a hIgG1-LALA-delK constant region, which can be detected specifically with the Alexa 647 goat anti-human IgG, F(ab’)₂ antibody that is specific for detecting the Fab portion of human IgGs. Multiple images were acquired with a Zeiss LSM 800 confocal microscope and quantitated using ImageJ. Care was taken to ensure that the signal intensities were not saturating. Approximately 10-20 NMJs were imaged and quantitated (individual dots/image) from each image. Binding of ARGX-119 at the NMJ was measured by calculating the ratio of the signal intensity for ARGX- 119 binding to the signal intensity for AChR expression.

*Post-hoc statistical analysis.* The obtained datasets (AChR signal intensity, ARGX-119 signal intensity and the ratio of ARGX-119 to AChR signal intensity) were independently analyzed by a biostatistician. For analysis, the background staining per image was subtracted from all signal intensities. The AChR density normalized by the mean AChR density of the vehicle group and the ratio of the ARGX- 119/AChR densities were statistically analyzed using the ‘mixed’ procedure in SAS Viya release V.03.04. The final models were run with restricted maximum likelihood. The assumptions of normality and homoscedasticity of the residuals with expectation zero were confirmed. The linear mixed model for the AChR normalized density included dose as a continuous fixed effect and image nested within dose as a random effect. The linear mixed model for the ratio of the ARGX-119/AChR densities included dose as a categorical fixed effect and image nested within dose as a random effect. Due to large differences in variability of the ratios between dosing groups, this model also incorporated heterogeneity in the residual variability between dosing groups. All dosing groups were pairwise compared to evaluate the effect of dose. Pairwise hypothesis tests were performed and p-values were corrected by applying Hommel’s procedure for multiple testing to have a family-wise error rate of 5%. The denominator degrees of freedom for the hypothesis tests were approximated [35]. In order to predict the dose of ARGX-119 that saturated MuSK, an Emax dose-response model (ARGX-119:AChR) was fitted to these ratios using the ‘MCPMod’ package in R v4.0.1. The Emax model was preferred above a sigEmax model as the predictions were very similar for both models, though the Emax model was selected for reporting due to its simplicity compared to a sigEmax model. The Emax model is characterized by three parameters according to the following equation:

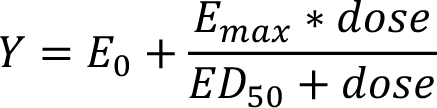

with

- E_0_ = predicted baseline/vehicle response
- E_max_ = asymptotic maximum change from vehicle effect
- ED_50_ = dose giving half of the asymptotic maximum effect

### Pharmacokinetic study in NHPs, rats and mice

These non-good laboratory practice studies where performed at animal facilities in Europe and were compatible with good laboratory practice regulations specified by regulatory authorities throughout the European Community, United States and Japan. Single doses of ARGX-119, ranging from 0.025 up to 200 mg/kg where tested in NHP (cynomolgus monkey), rats or mice. Serum was collected at several time points pre- and postdosing, and day of injection is referred as day 0. Three NHPs (2 males, 1 female) per group were intravenously injected, four male rats (low-dose range up to 0.5 mg/kg) per group and 6 rats (3 males, 3 female, high dose range from 0.5 mg/kg to 200 mg/kg) per group were intravenously injected, and 4 to 6 mice per group were intraperitoneally injected (4 females at 0.3 mg/kg, 4 males at 1 mg/kg, 2 males and 2 females at 10 mg/kg and 6 males at 20 mg/kg).

### Mouse model for *Dok7* CM

The *Dok7* CM mouse model was first described in [17], and experiments described here are performed by the same person with the same methods, using the same mouse strain (males and females). Isotype Ab used was Mota-hIgG1-LALAdelK. No statistical method was used to predetermine sample size. No data were excluded from the analyses. The experiments were not randomized. The investigators were not blinded to the genotype of the mice with the exception of the motor performance experiments.

## Acknowledgements

We thank Kathleen Moens, Rani Coppejans, Lieselot De Clercq and Leah Santana for experimental execution support, Ariëlla Van de Sompel for support with biostatistical analysis and Erwin Pannecoucke for contributing to the ITC experiments. Prof. Dr. Jan Verschuuren and Prof. Dr. Silvère van der Maarel for the vibrant discussions. The LUMC forms part of the European Reference Network for Rare Neuromuscular Diseases (ERN EURO-NMD) and the Netherlands Neuromuscular Disorders Center (NL-NMD). MGH receives support from a 2019 ZonMW VENI grant, LUMC Gisela © Fellowship 2021 and the 2023 Dutch incentive grant.

## Funding

This study was funded by argenx.

## Author contributions

R.V., J.O., B.V. and S.B. supervised, assisted in the design and interpretation of experiments and drafted the paper; C.S conducted antibody selections utilizing the SIMPLE Antibody™ platform, performed experiments, interpreted the data, contributed analysis tools and critically revised the paper. S.J., D.V., C.K., J.L., J.P. and R.A. performed experiments and interpreted the data. S.H., C.B., P.U., M.H and K.S supervised and assisted in the design and interpretation of experiments and critically revised the paper.

## Conflict of interest

R.V., B.V., C.S., C.B., P.U. and K.S. are (former) employees / consultants of argenx BV and are holders of employee equity in argenx.

J.O., S.K., S.B. are co-inventors on MuSK-related (pending) patents. S.B. is a consultant for argenx.

S.K. and S.B. have received research funding from argenx.

M.H. and J.P. are co-inventors on MuSK-related pending patents and receive royalties. LUMC receives royalties on a MuSK ELISA. M.H. is consultant for argenx.

## Issued patents

Granted US patent US 9,329,182 titled METHOD OF TREATING MOTOR NEURON DISEASE WITH AN ANTIBODY THAT AGONZES MUSK, by applicant New York University

The pending EP and US patent applications, derived from the international patent publication nr WO2020/055241 titled MUSK ACTIVATION, by applicant Academisch Ziekenhuis Leiden (h.o.d.n. LUMC)

The pending EP and US patent applications, derived from international patent publication nr WO2021/180676 titled MUSK AGONIST ANTIBODY, by co-applicants argenx BV and Academisch Ziekenhuis Leiden

Granted US patent US 11,492,401 titled THERAPEUTIC MUSK ANTIBODIES, by co-applicants New York University and argenx BV, with pending counterparts in multiple jurisdictions

The pending patent family with international patent publication nr WO2023/147489 titled ANTI- MUSK ANTIBODIES FOR USE IN TREATING NEUROMUSCULAR DISORDERS, by co-applicants argenx BV, Université de Montréal and New York University

The pending patent family with international patent publication nr WO2023/218099 titled IN UTERO TREATMENT OF A FETUS HAVING GENETIC DISEASE / NEUROMUSCULAR DISEASE, by applicant argenx BV

## Data and materials availability

All data are available in the manuscript or supplemental materials. The materials described herein may be available for research purposes. Any transfer of materials would be subject to a material transfer agreement.

## Notes

### Competing Interest Statement

C.S., B.V., K.S., C.K., J.L., C.B., P.U., K.S. and R.V. are employees of and have equity ownership in argenx BV.
SJB recieved financial support for research from argenx BV.

