## Supplemental Methods, Supplemental Figures 1-7, Supplemental Tables 1-5 for "ARGX-119, a therapeutic agonist antibody targeting MuSK"

### Supplemental Materials

#### Supplemental Methods

##### Serum samples, reagents

The human, NHP, mouse and rat MuSK protein sequences were based on the public available UniProtKB sequences with respective accession numbers O15146, A0A2K5USG2, Q61006 and Q62838 (UniProt). To confirm that these sequences are relevant for the selected species to be used during pre-clinical development of ARGX-119, cDNA sequencing was performed on cDNA obtained from humans, cynomolgus monkeys, BALB/c mice and Wistar rats. The obtained MuSK sequences showed 100% identity to the public sequences and were ordered for production (ImmunoPrecise Antibodies). 3F6c-hIgG1-LALAdelK and 13-3B5-hIgG4 are monoclonal antibodies targeting the MuSK Ig-like 1 domain on non-overlapping epitopes and are used in this study as assay tools or controls. Their variable domains were isolated from MuSK MG patients recruited in the MG outpatient clinic at the Leiden University Medical Centre (LUMC) as described previously [1]. 3F6c-hIgG1-LALAdelk was produced and purified at Immunoprecise Antibodies. 3B5-hIgG4 was produced and purified at Evitria SA (Zurich, Switzerland). To obtain an isotype control for ARGX-119, the variable domains of Motavizumab$^{TM}$, a humanized monoclonal antibody for the prevention of respiratory syncytial virus infection in high-risk infants, was formatted into the hIgG1-LALAdelK backbone and named 'Mota-hIgG1-LALAdelK'.

##### Growth and differentiation of myoblast cell lines

C2C12 mouse skeletal myoblasts, purchased from and authenticated by ECACC (CAT 91031101), were grown at 37 °C in growth medium (GM): Dulbecco's modified Eagle's medium (DMEM) containing high glucose (4.5 g/L), GlutaMAX$^{TM}$ (L-alanyl-L-glutamine) and sodium pyruvate (ThermoFisher Scientific), supplemented with 10% fetal bovine serum (FBS; Sigma Aldrich). Myoblast fusion and myotube differentiation were induced when myoblasts were 70% confluent by switching to differentiation medium (DM): DMEM high glucose, GlutaMAX™, pyruvate, supplemented with 1% heat-inactivated horse serum (Sigma Aldrich). Formation of C2C12 mouse myotubes took an average of five days. L6 rat skeletal myoblasts, purchased from and authenticated by ATCC (CAT CRL-1458), were grown at 37 °C in GM: DMEM, high glucose, GlutaMAX™, pyruvate, supplemented with 10% FBS. Myoblast fusion and myotube differentiation were induced when myoblasts were 70% confluent by switching to DM: DMEM, high glucose, GlutaMAX™, pyruvate, supplemented with 1% heat-inactivated horse serum and 0.5x Insulin-Transferrin-Selenium (100x ITS-G, ThermoFisher Scientific). Formation of L6 rat myotubes took an average of six days. CynoSMM primary cynomolgus monkey skeletal muscle myoblasts, purchased from and authenticated by Cell Biologics (CAT MK-6167), were grown at 37 °C in GM: Complete Smooth Muscle Cell Medium (Cell Biologics), completed with growth factor supplement kit containing FBS, epidermal growth factor (EGF), fibroblast growth factor (FGF), hydrocortisone, insulin and antibiotic-antimycotic solution (Cell Biologics). Subculture plates were coated with 0.1% porcine gelatin (Sigma Aldrich). Gelatin-coating was not performed on assay plates, as the gelatin interferes with the *in vitro* read-outs. Myoblast fusion and myotube differentiation were induced when myoblasts were 90% confluent by switching to DM: DMEM, high glucose, GlutaMAX™, pyruvate, supplemented with 1% heat-inactivated horse serum and 1x ITS-G. Formation of CynoSMM primary cynomolgus monkey myotubes took an average of five days. MB135 human skeletal muscle myoblasts immortalized with hTERT and CDK4, licensed from Professor Stephen J. Tapscott (Division of Basic Sciences, Fred Hutchinson Cancer Research Center, Seattle, WA, USA), were grown at 37 °C in GM: Ham's F-10 Nutrient Mix containing L-glutamine (ThermoFisher Scientific), supplemented with 15% FBS, 10 ng/mL human recombinant FGF (rhFGF, Promega), and 1 µM dexamethasone (Sigma Aldrich). Myoblast fusion and myotube differentiation were induced when myoblasts were 90% confluent by switching to DM: DMEM, high glucose, GlutaMAX™, pyruvate, supplemented with 1% heat-inactivated horse serum and 1x ITS-G. Formation of MB135 human myotubes took an average of four days. HSMM primary human skeletal muscle myoblasts, purchased from and authenticated by Lonza (via Clonetics, CAT CC-2580), were grown at 37 °C in GM: Skeletal Muscle Cell Growth Basal

Medium (SkGMTM, Lonza), completed with SKGM™ SingleQuots™ supplements and growth Factors containing FBS, human EGF (hEGF), dexamethasone, L-glutamine, and gentamicin/amphotericin-B (GA) (Lonza). Myoblast fusion and myotube differentiation were induced when myoblasts were 90% confluent by switching to DM: DMEM high glucose, GlutaMAX$^{TM}$, pyruvate, supplemented with 1% heat-inactivated horse serum. Formation of HSMM primary human myotubes took an average of four days followed by 1 day back on GM to improve cell viability. A maximum of ten passages were maintained for all the myoblast cell lines before starting a new culture.

**C1q ELISA**

Recombinant human MuSK-ECD was coated overnight at 4°C on a MaxiSorp plate (Thermo Scientific). The next day the assay plate was blocked with 1% casein blocking buffer (Bio-Rad) for 1 hr at RT. ARGX-119 and Fc variants (WT, LALAPG) were prepared fresh by diluting the stock solution in assay buffer (0.1% Casein in 1xPBS). These antibody dilutions were added in duplicate to the assay plate and incubated for 1 hr at RT. Next, human serum (in-house pool lot RB-2022-0027_pool05), NHP serum (cynomolgus monkey, in-house pool lot B20240103), mouse serum (C57BL/6, MSE492316, BioIVT) and rat serum (Sprague Dawley, ab7488, Abcam) were diluted to 10% in assay buffer. The serum dilutions were incubated on the plate for 1hr at RT. C1q was detected by incubation of a biotinylated anti-C1q antibody (MA1-40312, Invitrogen) for 1h at RT followed by incubation with streptavidin-conjugated-HRP for 1hr at RT. TMB substrate was added, color development reaction was terminated by adding sulfuric acid, and OD 450nm was measured on an Infinite M Nano plate reader (Tecan). Results were processed using Graphpad Prism 9 software.

**Supplemental Figures and Figure Legends**

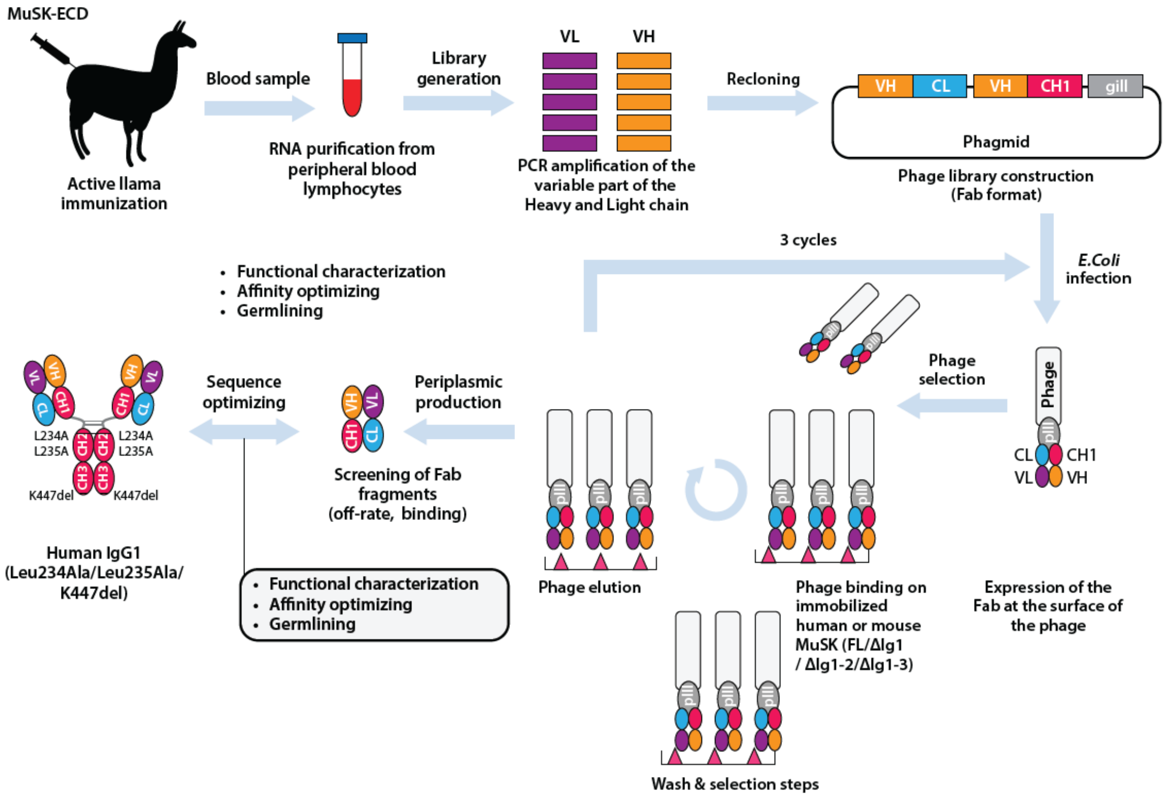


**Supplemental Figure 1: Generation of ARGX-119 using SIMPLE Ab platform**

Graphical representation of the SIMPLE Ab platform, from llama immunizations with human MuSK ECD, to phage library constructions, phase display selections, functional characterization and germlining to the production of the final lead mAb.

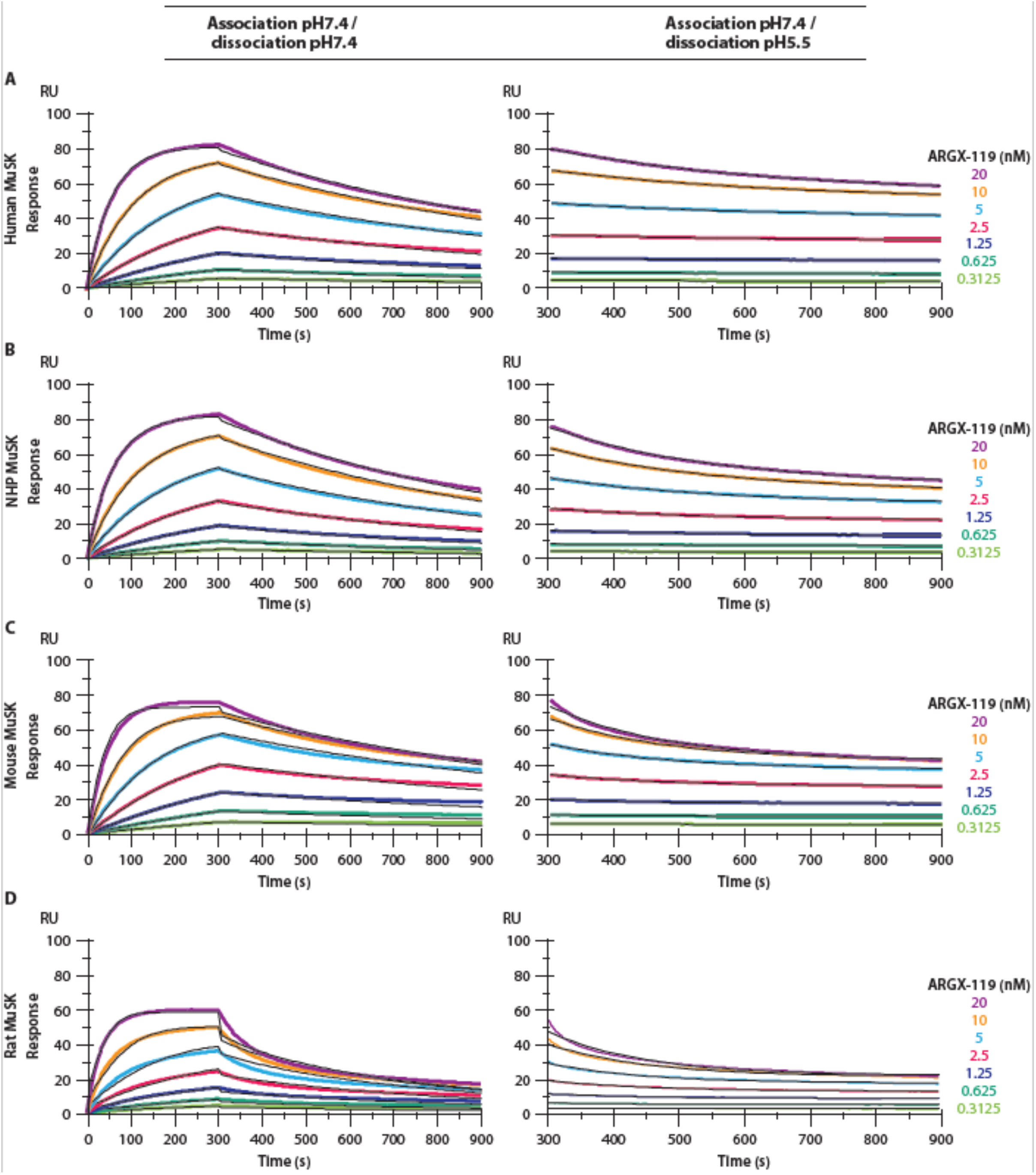


**Supplemental Figure 2: SPR titrations of ARGX-119 binding to MuSK**

SPR binding of ARGX-119 to MuSK. CM5 chip was coated with anti-human IgG-Fc and ARGX-119 was captured at low densities. Sensorgrams show the binding kinetics of the titrations of the different MuSK proteins using a dissociation pH of 7.4 or 5.5. Kinetic parameters at pH7.4 were calculated using a Langmuir 1:1 interaction model. Dissociation rates at pH 5.5 were calculated using the T200 software (v3.2) dissociation fit model.

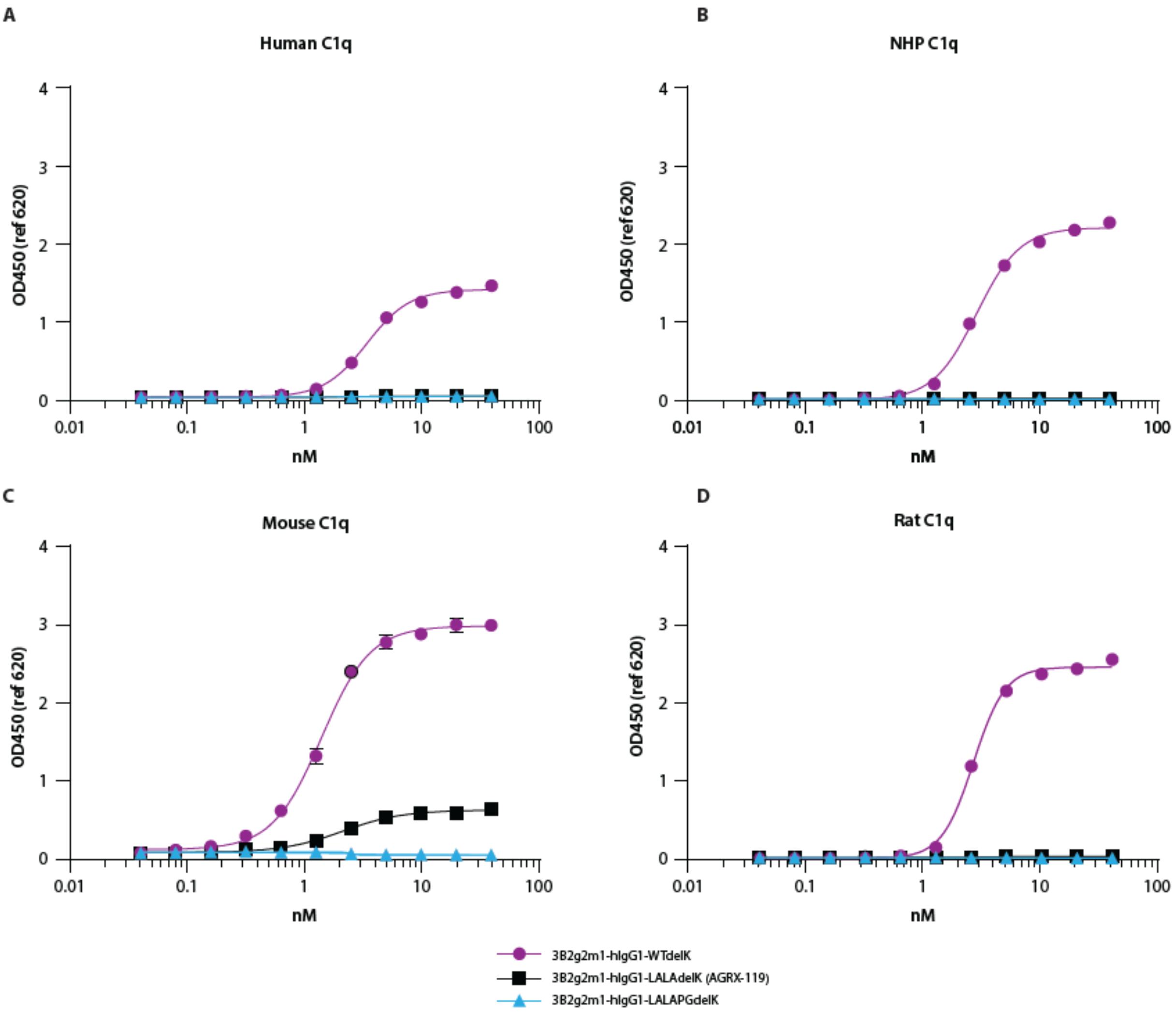


**Supplemental Figure 3: ARGX-119 has diminished reactivity to C1q**

ELISA to test C1q reactivity. MuSK was coated and mAbs to MuSK were allowed to bind. Next, serum (containing C1q) was incubated and signal was generated via anti-C1q detection. Serum containing C1q from human, NHP, mouse and rat were tested for reactivity with ARGX-119 and controls (WT and LALAPG Fc variants).

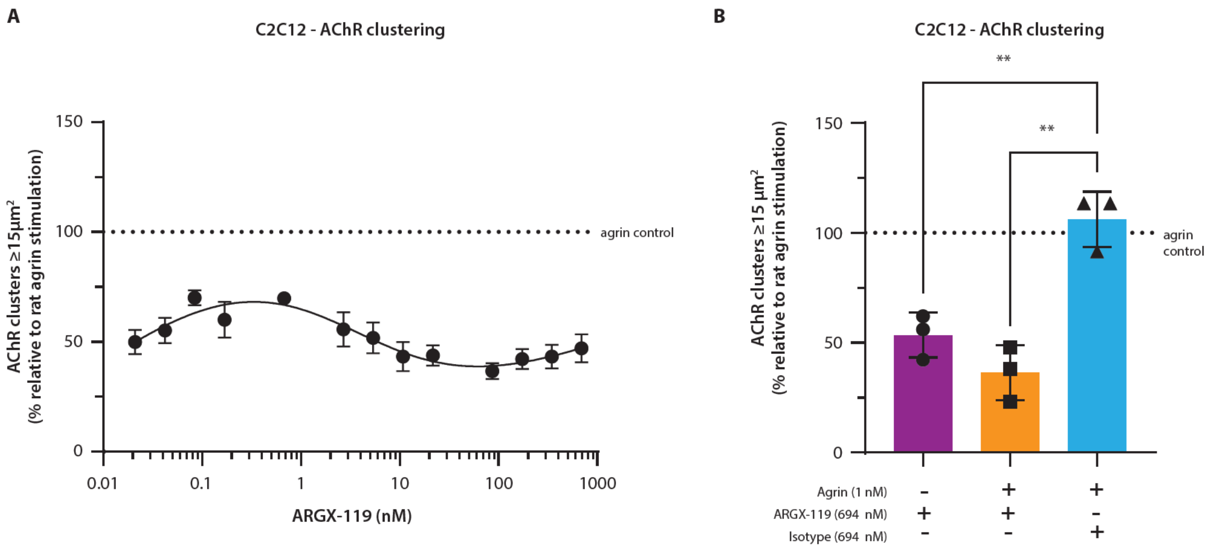


**Supplemental Figure 4: High dose ARGX-119 data on ≥15 µm² AChR clusters**

(A) AChR clusters ≥ 15 µm$^2$ in mouse C2C12 myotubes treated for 24 hr with a dose-range of ARGX-119. (B) AChR clusters ≥ 15 µm$^2$ in mouse C2C12 myotubes treated for 24 hr with Agrin, ARGX-119, isotype control mAb, or in combination. One-way ANOVA with Dunnet's multiple comparison (** $p<0.01$). AChR clustering data were generated from 3 individual experiments (n=3), each done in quadruplicate.

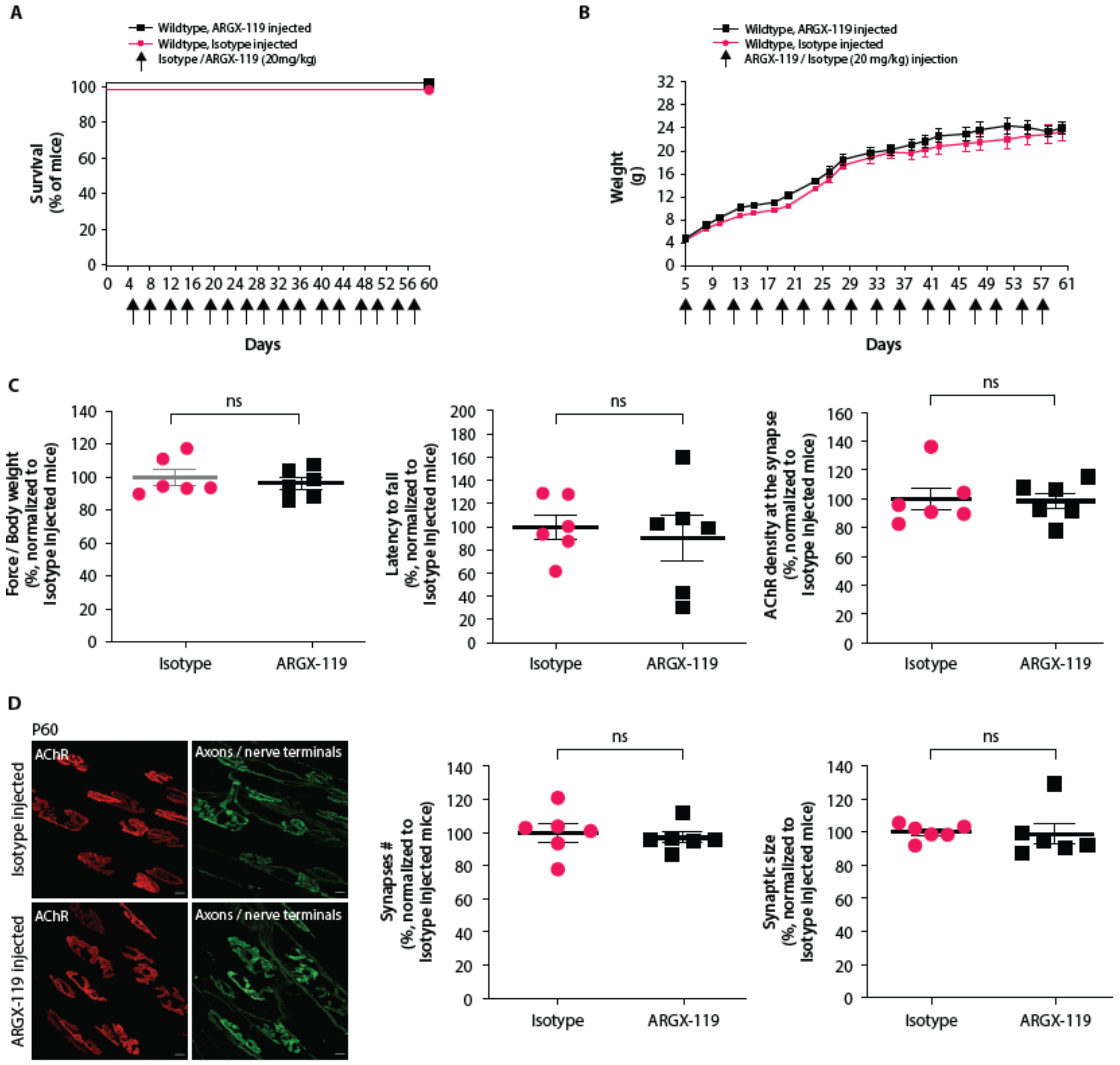


**Supplemental Figure 5: ARGX-119 dosing in wild-type mice**

Wild-type mice (C57BL/6) were injected twice a week with either Isotype (20 mg/kg, n=6) or ARGX-119 (20 mg/kg, n=6) starting at P4 until P60. (A) Plots show the percentage of mice surviving for each treatment over 60 days. Log-rank (Mantel-Cox) test. (B) Wild-type mice injected twice a week with ARGX-119 gained weight, like wild-type mice treated with Isotype. Plots show averaged data points and mean ± SEM. (C) Motor performance of wild-type mice injected twice a week with ARGX-119, as assessed by grip strength and the latency to fall from a rotarod at P60, was similar to Isotype-injected wild-type mice. The scatter plots show the values for 6 mice per group and the mean ± SEM. Percentage values were normalized to Isotype-injected mice. Two-sided Student's *t*-test (ns, not significant). (D) Left, diaphragm muscles from P60 wild-type mice injected with Isotype or ARGX-119 were stained to label AChRs (red) as well as motor axons and nerve terminals (green). Scale bar = 10 µm. Right, synapse number, size or AChR density (>50 synapses per diaphragm muscle from 6 mice in each group). Scatter plots show the values in percentage for 6 mice per group and the mean ± SEM. Percentage values were normalized to isotype-injected mice. Two-sided Student's *t*-test (ns, not significant).

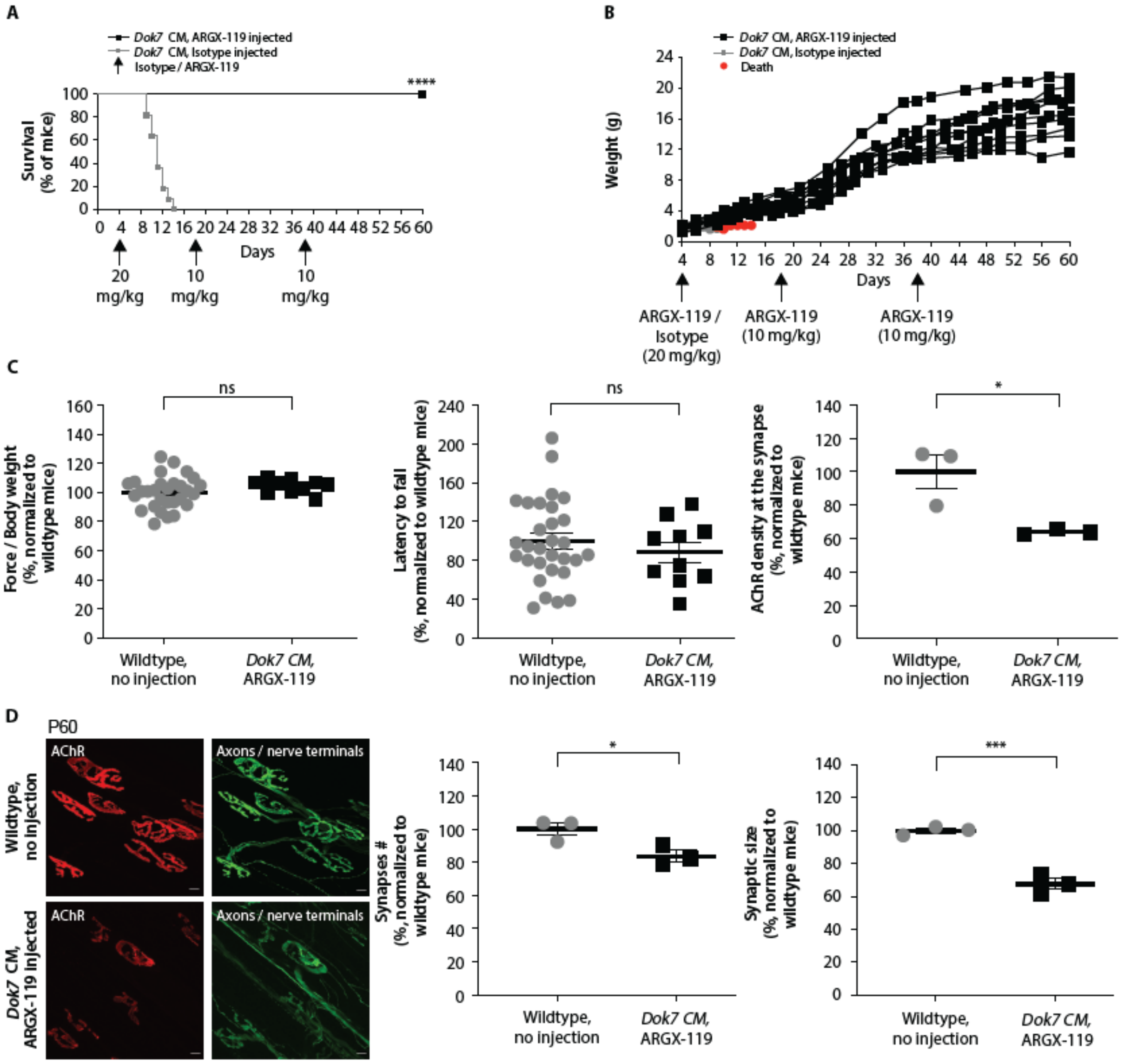


**Supplemental Figure 6: ARGX-119 restores synaptic differentiation and motor behavior and prevents early postnatal lethality in *Dok7* CM mice.**

(A) *Dok7* CM mice (C57BL/6-CBA mixed background) survive for two weeks postnatally. *Dok7* CM mice were injected with ARGX-119 (n=10) at P4 (20 mg/kg), P18 (10 mg/kg) and P38 (10 mg/kg), and all of these mice survived until the end of experiment, when mice were P60. Mice (n = 11) injected with isotype mAb died within two weeks after birth. Plots show the percentage of mice surviving for each treatment over 60 days. Log-rank (Mantel-Cox) test (p, ****<0.00005). (B) Chronic injection with ARGX-119 restored weight gain in *Dok7* CM mice, unlike *Dok7* CM mice treated with isotype control mAb. Plots show the values for individual mice over a period of 60 days. (C) Motor performance of

*Dok7* CM mice, as assessed by grip strength and the latency to fall from a rotarod at P60, were fully restored by treatment with ARGX-119. The scatter plots show the values for 30 wild-type mice and 10 *Dok7* CM mice rescued with ARGX-119 and the mean in percentage values (mean ± SEM), normalized to wild-type mice. Two-sided Student's t-test (ns, not significant). (D) Diaphragm muscles from P60 mice were stained with Alexa 488-α-BGT to label AChRs (red) and motor axons/nerve terminals (green). In *Dok7* CM mice treated with ARGX-119, synapses matured from a simple, plaque-like shape to a complex, pretzel-like shape, characteristic of mature murine neuromuscular synapses. Scale bar = 10 µm. Chronic injection with ARGX-119 restored the number and size of synapses and the density of synaptic AChRs in *Dok7* CM mice. (>50 synapses per diaphragm muscle from 6 mice in each group). Scatter plots show the values (mean ± SEM) for 3 wild-type mice and 3 *Dok7* CM mice injected with ARGX-119. Percentage values were normalized to wild-type mice average. Two-sided Student's t-test (p, $*<0.05$, $***<0.0005$).

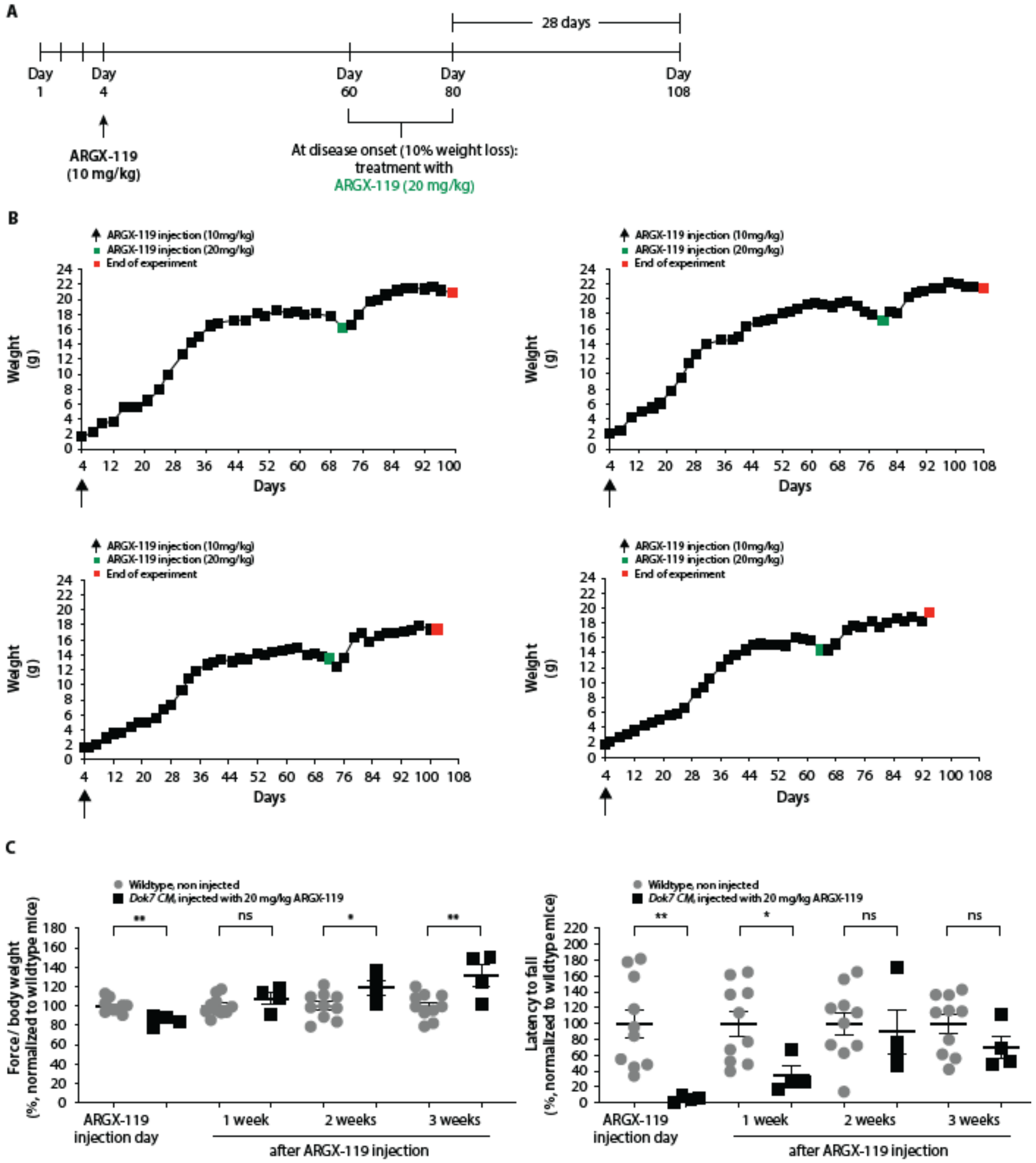


**Supplemental Figure 7: 20 mg/kg ARGX-119 fully rescues relapsed *Dok7* CM mice**

(A) Strategy for testing potential of ARGX-119 to reverse disease relapse in adult *Dok7* CM mice. We injected *Dok7* CM mice (C57BL/6-CBA background) with ARGX-119 (10 mg/kg) at P4 and then discontinued antibody treatment. When mice began to lose weight (~10% weight loss), they were re-injected with ARGX-119 (20 mg/kg), and monitored for 4 weeks. (B) *Dok7* CM mice injected with a single dose of ARGX-119 (10 mg/kg) at P4 ultimately began to lose weight. When weight loss was reduced by 10%, *Dok7* CM mice were reinjected with 20 mg/kg ARGX-119. The weight loss of *Dok7* CM mice was reversed within one week after restarting ARGX-119 treatment. Red dots indicate mice that were sacrificed at the end of the experiment. (C) *Dok7* CM mice injected with a single dose of ARGX-119 (10 mg/kg) at P4 displayed motor deficits at disease onset as assessed by grip strength and the latency to fall from a rotarod. The motor skills of the recovered *Dok7* CM mice were similar to those of non-injected wild-type mice. Plots show individual data points and mean (± SEM) in percentage normalized to non-injected mice. Two-sided Student's t-test (ns, non-significant, p, *<0.05, **<0.005, ****<0.00005).

**Supplemental Tables**

Supplemental Table 1: ARGX-119 affinity measurements across species

| | | SPR | | | | | | ITC | | | | |
|---|---|---|---|---|---|---|---|---|---|---|---|---|
| MuSK-ECD | Dissociation pH | AVG capture Level (RU) | %CV | Ka (1/Ms) | kd (1/s) | KD (nM) | U-value | KD (nM) | ΔH (kcal/mol) | -TΔS (kcal/mol) | ΔG (kcal/mol) | n |
| Human | 7.4 | 95 | 1.4 | 9.0E+05 | 1.0E-03 | 1.2 | 1 | 1.6 ± 2.7 | -35.7 ± 6.5 | 23.2 ± 6.8 | 12.6 ± 0.3 | 4.3 ± 1.7E-03 |
| | 5.5 | 82 | 3.9 | n/a | 2.8E-03 | n/a | n/a | n/a | n/a | n/a | n/a | n/a |
| Nonhuman primates | 7.4 | 91 | 0.7 | 1.7E+06 | 1,0E-03 | 0.6 | 1 | 2.2 ± 4.5 | -24.6 | 12.3 | -12.3 | 0.6 ± 2.9E-03 |
| | 5.5 | 78 | 3.1 | n/a | 4.0E-03 | n/a | n/a | n/a | n/a | n/a | n/a | n/a |
| Mouse | 7.4 | 93 | 0.6 | 7.6E+05 | 1.3E-03 | 1.7 | 1 | 13.1 ± 1.6 | -32.9 ± 8.3 | 20.7 ± 8.6 | -12.2 ± 0.4 | 0.2 ± 2.2E-03 |
| | 5.5 | 79 | 2.4 | n/a | 5.5E-03 | n/a | n/a | n/a | n/a | n/a | n/a | n/a |
| Rat | 7.4 | 90 | 2.5 | 2.3E+06 | 3.4E-03 | 1.5 | 3 | 5.4 ± 1.6 | -37.2 ± 0.9 | 25.5 ± 1.0 | -11.8 ± 0.1 | 0.3 ± 3.9E-03 |
| | 5.5 | 77 | 1.9 | n/a | 6.8E-03 | n/a | n/a | n/a | n/a | n/a | n/a | n/a |

n/a – not applicable (different buffers used for the association and dissociation phase)

Supplemental Table 2: ARGX-119 binding to Fc gamma receptors

| | | Averaged binding KD (nM), N=3 | | |
|---|---|---|---|---|
| **Fc gamma receptor** | | 3B2g2m1-hIgG1-WTdelK | 3B2g2m1-hIgG1-LALAdelK (ARGX-119) | 3B2g2m1-hIgG1-LALAPGdelK |
| Human | hFcγRI | 6.5 | 126.6 | Low binding (>5000) |
| Human | hFcγRIIa 131H | 780.0 | Low binding (>5000)* | ND |
| Human | hFcγRIIa 131R | 1080.2 | Low binding (>5000) | ND |
| Human | hFcγRIIb | Low binding (>5000) | ND | ND |
| Human | hFcγRIIIa 158F | 1635.9 | Low binding (>5000) | ND |
| Human | hFcγRIIIa 158V | 344.8 | Low binding (>5000) | ND |
| Human | hFcγRIIIb NA1 | >5000 | ND | ND |
| Human | hFcγRIIIb NA2 | >5000 | ND | ND |
| Non-human primate | cFcγRIIa | 2074.6 | ND | ND |
| Non-human primate | cFcγRIIb | 1422.7 | Low binding (>5000) | ND |
| Non-human primate | cFcγRIII | 127.6 | Low binding (>5000) | ND |
| Mouse | mFcγRI | 69.2 | Low binding (>5000) | ND |
| Mouse | mFcγRIIb | 814.2 | ND | ND |
| Mouse | mFcγRIII | 1028.7 | ND | ND |
| Mouse | mFcγRIV | 392.2 | Low binding (>5000) | ND |
| Rat | rFcγRI | 316.5 | Low binding (>5000) | Low binding (>5000) |
| Rat | rFcγRIIa | ND | ND | ND |
| Rat | rFcγRIIb | 4960.4 | Low binding (>5000) | ND |
| Rat | rFcγRIII | 470.9 | Low binding (>5000) | ND |

ND: not detected

Supplemental Table 3: ARGX-119 MuSK phosphorylation and AChR clustering dose response values

| | Myotubes | Analytical runs | ARGX-119 (nM) | | | | | |
|---|---|---|---|---|---|---|---|---|
| | | | **EC5** | **EC10** | **EC25** | **EC50** | **EC75** | **EC95** |
| MuSK phosphorylation | Human (immortalized) – MB135 | N=3 | 0.09 | 0.12 | 0.18 | 0.26 | 0.39 | 0.76 |
| | Human (primary) – HSMM | N=3 | 0.05 | 0.07 | 0.13 | 0.23 | 0.41 | 1.11 |
| | Rat (immortalized) – L6 | N=3 | 0.32 | 0.44 | 0.72 | 1.20 | 2.04 | 5.40 |
| | Mouse (immortalized) – C2C12 | N=3 | 0.22 | 0.31 | 0.51 | 0.85 | 1.41 | 3.27 |
| AChR clustering (≥ 3 µm²) | Human (immortalized) – MB135 | N=3 | 0.00015 | 0.00034 | 0.00116 | 0.00391 | 0.01324 | 0.10283 |
| | Mouse (immortalized) – C2C12 | N=3 | 0.00100 | 0.00300 | 0.00700 | 0.01900 | 0.05700 | 0.37700 |

Supplemental Table 4: *In vitro* off target screening

| Sample ID | Gene ID | Protein Name | Accession | Library screen (fixed cells) | | Confirmation screen (fixed cells) | | Confirmation screen (live cells) | Comments |
|---|---|---|---|---|---|---|---|---|---|
| | | | | Rep 1 | Rep 2 | Rep 1 | Rep 2 | | |
| ARGX-119 | MUSK | Muscle-specific kinase | NM_005592.3 | n/a | n/a | medium/strong | medium/strong | medium/strong | 869aa, isoform 1, single-pass type I membrane protein |
| X17 | MuSK | Muscle-specific kinase | NM_005592.3 | strong | strong | medium/strong | medium/strong | medium/strong | 869aa, isoform 1, single-pass type I membrane protein |
| | EPHB1 | Ephrin type-B receptor 1 | NM_004441.4 | very weak | weak | weak | very weak | very weak | 984aa, isoform 1, single-pass type I membrane protein |
| | EPHB2 | Ephrin type-B receptor 2 | NM_004442.6 | weak/medium | medium | medium | medium | weak/medium | 987aa, isoform 1, single-pass type I membrane protein |

n/a – not applicable (not tested in library screen), aa – amino acid

Supplemental Table 5: ARGX-119 PK parameters in NHPs, rats and mice

| | Dose | Cmax | AUCINF | AUCINF_D | t1/2,z | Cl | Vz |
|---|---|---|---|---|---|---|---|
| | mg/kg | µg/mL | day*µg/mL | (day*µg/mL)/ (mg/kg) | day | mL/day/kg | mL/kg |
| NHP | | | | | | | |
| Study 1 | 0.1 | 2.8 | 4.4 | 44 | 12 | 23 | 410 |
| | 0.5 | 14 | 55 | 110 | 10 | 9.1 | 130 |
| | 2 | 60 | 439 | 219 | 9.5 | 4.6 | 62 |
| | 10 | 311 | 2726 | 273 | 14 | 3.7 | 73 |
| Rat | | | | | | | |
| Study 2 | 0.025 | 0.78 | 3 | 120 | 10.5 | 8.9 | 126 |
| | 0.05 | 1.51 | 7.44 | 149 | 10.9 | 6.7 | 106 |
| | 0.1 | 3.4 | 14.1 | 141 | 8.8 | 7.2 | 90.7 |
| | 0.5 | 17.5 | 101 | 202 | 11.9 | 5 | 85.1 |
| Study 1 | 0.5 | 16.2 | 92.7 | 185 | 10.8 | 5.5 | 85.1 |
| | 5 | 141 | 967 | 193 | 13 | 5.2 | 97.2 |
| | 50 | 1610 | 11100 | 223 | 12.2 | 4.5 | 78.6 |
| | 200 | 6680 | 40700 | 204 | 13.3 | 5 | 95.6 |
| Mouse | | | | | | | |
| Study 1 | 0.25 | 8.3 | 47 | 186 | 7 | 5.4 | 54 |
| | 2.5 | 110 | 1072 | 429 | 12 | 2.3 | 39 |
| | 25 | 810 | 6887 | 275 | 14 | 3.6 | 72 |
| | 100 | 3370 | 37434 | 374 | 17 | 2.7 | 64 |
